# Membrane-associated periodic skeleton regulates major forms of endocytosis in neurons through a signaling-driven positive feedback loop

**DOI:** 10.64898/2025.12.12.693977

**Authors:** Jinyu Fei, Yuanmin Zheng, Caden LaLonde, Yuan Tao, Ruobo Zhou

## Abstract

Endocytosis is an evolutionarily conserved process that enables neurons to internalize signaling receptors and membrane proteins, maintaining cellular homeostasis and supporting rapid responses to extracellular cues. The neuronal membrane-associated periodic skeleton (MPS), a lattice-like cytoskeletal structure composed of actin and spectrin, has been shown to restrict clathrin-mediated endocytosis (CME) at the axon initial segment (AIS) of neurons, by gating clathrin-coated pit (CCP) formation through membrane-localized “clearing” structures that are devoid of MPS. However, the extent to which the MPS regulates diverse forms of endocytosis across neuronal compartments, and how it is dynamically remodeled to permit trafficking on demand, remain unknown. While CME is relatively well characterized in neurons, the subcellular localization and physiological relevance of caveolin-mediated endocytosis, flotillin-mediated endocytosis, and fast endophilin-mediated endocytosis (FEME) have remained largely unclear. Here, we show that all four major endocytic pathways—CME, caveolin-, flotillin-, and FEME— are spatially gated by the MPS and occur specifically within MPS-free “clearing” zones distributed across both axonal and somatodendritic compartments of mature neurons. These results, for the first time, map the spatial landscape of these lesser-understood pathways in neurons and reveal a unifying principle of cytoskeletal gating across endocytic mechanisms. Disruption of the MPS markedly enhances both basal and ligand-induced endocytosis across all four pathways, establishing its broad inhibitory role in pit initiation. We further discover that endocytosis can, in turn, remodel the MPS through a novel signaling-driven feedback loop: ligand-triggered endocytosis activates ERK signaling, which promotes calpain- and caspase-mediated spectrin cleavage. This targeted cytoskeletal degradation facilitates further rounds of endocytosis, forming a self-reinforcing circuit that couples membrane trafficking with cortical architecture remodeling. Finally, we show that the MPS limits amyloid precursor protein (APP) endocytosis and thereby suppresses amyloid-β 1-42 (Aβ42) production and neuronal apoptosis, implicating MPS integrity in the regulation of neurodegenerative processes such as Alzheimer’s disease. Together, our findings establish the MPS as a dynamic, signal-responsive modulator of endocytosis and neuronal health. This work uncovers a general spatial gating mechanism that applies to diverse endocytic pathways, introduces a cytoskeleton-centered feedback loop for signal-dependent remodeling, and expands the functional significance of the MPS from passive structural support to active regulation of neuronal homeostasis and disease susceptibility.

**Teaser:** Membrane skeleton gates endocytosis and remodels in response to signals, linking membrane dynamics to neuronal health and Alzheimer’s risk.

## Introduction

Endocytosis is a fundamental and evolutionarily conserved cellular process, essential for a wide range of organisms, including yeast, animals, and plants(*1*). It plays a critical role in maintaining the homeostasis of cell membrane constituents by recycling specific lipid and protein components of the plasma membrane and in facilitating the cell’s nutrient uptake and communication with its extracellular environment and neighboring cells by internalizing and sampling extracellular materials, such as fluids, biomolecules, and microbes. In neurons, endocytosis regulates nerve cell migration and adhesion during brain development, establishes neuronal differentiation and axon-dendrite polarity, and governs synaptic activity and plasticity by controlling synaptic transmission and shaping the size, structure, and number of synapses, thereby influencing brain development, functions, and activities such as learning and memory(*2*). Conversely, dysregulation of endocytosis has been implicated in the pathogenesis of various neurological disorders. Genetic studies have identified mutations which alter the expression of endocytic machinery genes in neurodegenerative diseases such as Alzheimer’s disease, Parkinson’s disease, and amyotrophic lateral sclerosis.

The regulation of endocytosis relies on the intricate interplay of protein-specific interactions, signaling pathway activation, and cytoskeleton rearrangement. While significant progress has been made in understanding the molecular mechanisms underlying endocytic pathways, many aspects of the molecular mechanisms underlying neuronal endocytosis remain poorly understood. To facilitate the uptake of diverse extracellular materials, various distinct endocytic mechanisms have been identified in non-neuronal cells, including clathrin-mediated endocytosis (CME, a clathrin- and dynamin-dependent pathway), lipid raft-mediated endocytosis (LRME, a clathrin-independent but dynamin-dependent pathway), fast endophilin-mediated endocytosis (FEME, a clathrin-independent but dynamin-dependent pathway for rapid ligand-driven internalization of specific membrane proteins), macropinocytosis, and phagocytosis. Among these, CME is the most well-studied endocytic pathway in neurons and has been shown to occur ubiquitously across nearly all neuronal subcompartments, including the soma, axon shaft, dendrite shaft, and both pre- and post-synaptic sites. In contrast, much less is known about the localization and regulation of other endocytic pathways in neurons. A recent study has reported a novel cell surface-localized structure in neurons, termed “clearings”, which regulates CME at the axon initial segment (AIS) and proximal axon segments near the AIS(*8*). These “clearings” are empty, membrane-associated holes surrounded by an actin-spectrin-based cytoskeletal network. Clathrin-coated pits (CCPs) preferentially form at the center of these pre-existing “clearings” and are stabilized as long-lived and stalled structures before undergoing scission and conclusive endocytosis in response to plasticity-inducing stimulation. Despite these insights, several critical questions remain unresolved. First, although the initial study reports that the “clearings” of the actin-spectrin scaffold exclusively formed at AIS and proximal axon next to AIS and were absent from soma and dendrites, this observation was made using relatively immature neurons at 14 days in vitro (DIV 14), when the actin-spectrin scaffold has not fully developed in the somatodendritic compartments. Our previous systematic investigation of the developmental progression of the actin-spectrin-based membrane-associated periodic skeleton (MPS) demonstrated that MPS formation occurs much slower in somatodendrites than in axons, with the MPS network density in soma and dendrites reaching saturation only around DIV 21-28(*9*). Therefore, it remains an open question whether similar CCP-containing “clearings” exist in distal axons or somatodendritic compartments in more mature (DIV 21 or older) neurons. Second, whether clathrin-independent endocytic pits, such as those involved in LRME or FEME, also preferentially localize to the centers of MPS-free clearings, and whether their endocytic activity is similarly regulated by the MPS, have not been investigated. Third, the mechanisms that control the formation, size, and spatial distribution of these clearings in response to increased endocytic demand, such as during neuronal stimulation, are largely unexplored. Given that the MPS inhibits ligand-induced CME, a critical unresolved question is how the MPS is locally disassembled, through the formation or expansion of clearings, to accommodate elevated endocytic activity during periods of rapid neuronal signaling.

Here, using super-resolution microscopy(*10, 11*) in combination with quantitative cell biology approaches, we systematically investigated four major types of endocytosis in neurons: CME, caveolin-mediated endocytosis and flotillin-mediated endocytosis(*12*) (two prominent forms of LRME), and FEME(*13*). Similar to CME, we found that caveolin-mediated endocytosis, flotillin-mediated endocytosis, and endophilin-mediated endocytosis (i.e., FEME) can occur broadly across nearly all neuronal subcompartments, including the soma, AIS, axon shaft, and dendrite shaft. Notably, we identified CCP-centered clearings not only in the proximal axon shaft as previously reported(*8*), but also in distal axons and somatodendritic regions of mature neurons. For the other three types of endocytic pathways, we identified analogous “clearings” formed by the actin-spectrin-based MPS, with caveolin-, flotillin-, or endophilin-enriched nanodomains located at their centers, resembling the previously reported CCP-centered clearings. Disruption of the neuronal MPS enhanced both basal endocytosis mediated by clathrin, caveolin, flotillin, and endophilin, as well as ligand-induced receptor endocytosis across these pathways. These findings suggest that the MPS serves as a physical barrier to suppress endocytosis for all the four endocytosis types. We further demonstrated that ligand-induced receptor endocytosis mediated by these proteins activates downstream ERK signaling cascades and proteases (i.e., calpains and caspases), which in turn can promote MPS disassembly (i.e., enlarge the total area of “clearings”), forming a positive feedback loop that further enhances endocytosis. These findings indicate that the MPS acts as a dynamically regulated barrier to modulate endocytosis rates. Finally, we demonstrated that the MPS exerts a neuroprotective role by negatively regulating amyloid precursor protein (APP) endocytosis through the same positive feedback loop. Given that APP endocytosis is upregulated during neuronal aging and in Alzheimer’s disease, this MPS-mediated regulation may serve as a protective mechanism against disease-associated endocytic dysregulation. Together, these findings provide structural and mechanistic insights into the regulation of diverse neuronal endocytosis pathways under both physiological and pathological conditions.

## Results

### Compartment-resolved mapping of four endocytic pit types reveals broad suppression of basal endocytosis by the neuronal MPS

As studies of neuronal endocytosis have primarily focused on CME and it remains largely unclear which neuronal compartments support other endocytic pathways, we first sought to develop immunofluorescence (IF)-based assays capable of accurately mapping the spatial distributions of four major types of endocytic pits, including the endocytic pits for CME (i.e., CCPs), caveolin-mediated endocytosis, flotillin-mediated endocytosis, and FEME, respectively, across different neuronal compartments.

While the IF staining protocol for visualizing CCPs in neurons has been well documented(*8*), staining protocols for other endocytic pits in neurons are underdeveloped. Previous studies have reported that the commonly used cell permeabilization method using Triton X-100 substantially alters the staining patterns of lipid-raft-mediated endocytic pits(*14*). To overcome this, we expressed moderate level of GFP-tagged caveolin-1(Cav1), flotillin-1 (Flot1), or endophilin-A2 (EndoA2) in cultured neurons through low-titer lentiviral transduction, and tested IF protocols previously developed for non-neuronal systems (**Fig. S1A, B, C**). Neurons were immunostained with IF-validated antibodies against endogenous Cav1, Flot1(*12*), and EndoA2(*13*), respectively. The observed strong colocalization between immunostained puncta and the puncta in the GFP channel, which are likely endocytic pits, confirmed the specificity of these antibodies and validated the IF-based assays for visualizing these endocytic pits in neuronal systems.

Using these IF-based assays, with immunostaining for endogenous clathrin, Cav1, Flot1, and EndoA2, and structured illumination microscopy (SIM), a super-resolution imaging method with a lateral resolution of 100-120 nm, we investigated the basal-level spatial distributions of the four major types of endocytic pits across distinct neuronal compartments, including the AIS, distal axons, soma, and dendrites. To accurately distinguish these neuronal compartments, we used neurofascin, tau, and MAP2 as markers for the AIS, distal axonal segments, and dendrites, respectively (**Fig. 1A, B, C, D and Fig. S2C, D, E, F**). We quantified basal-level densities of these endocytic pits using endocytic pit area fraction, defined as the ratio of total endocytic pit area to the corresponding compartmental area demarcated by these markers. We found that all four types of endocytic pits were present in all neuronal compartments. CCPs and Flot1-pits were denser in distal axons than in any other neuronal compartments, whereas Cav1- and EndoA2-pits were denser in dendrites compared to other neuronal compartments (**Fig. 1E**), indicating that these four endocytic pathways are differentially regulated across distinct neuronal compartments.

**Fig. 1.**
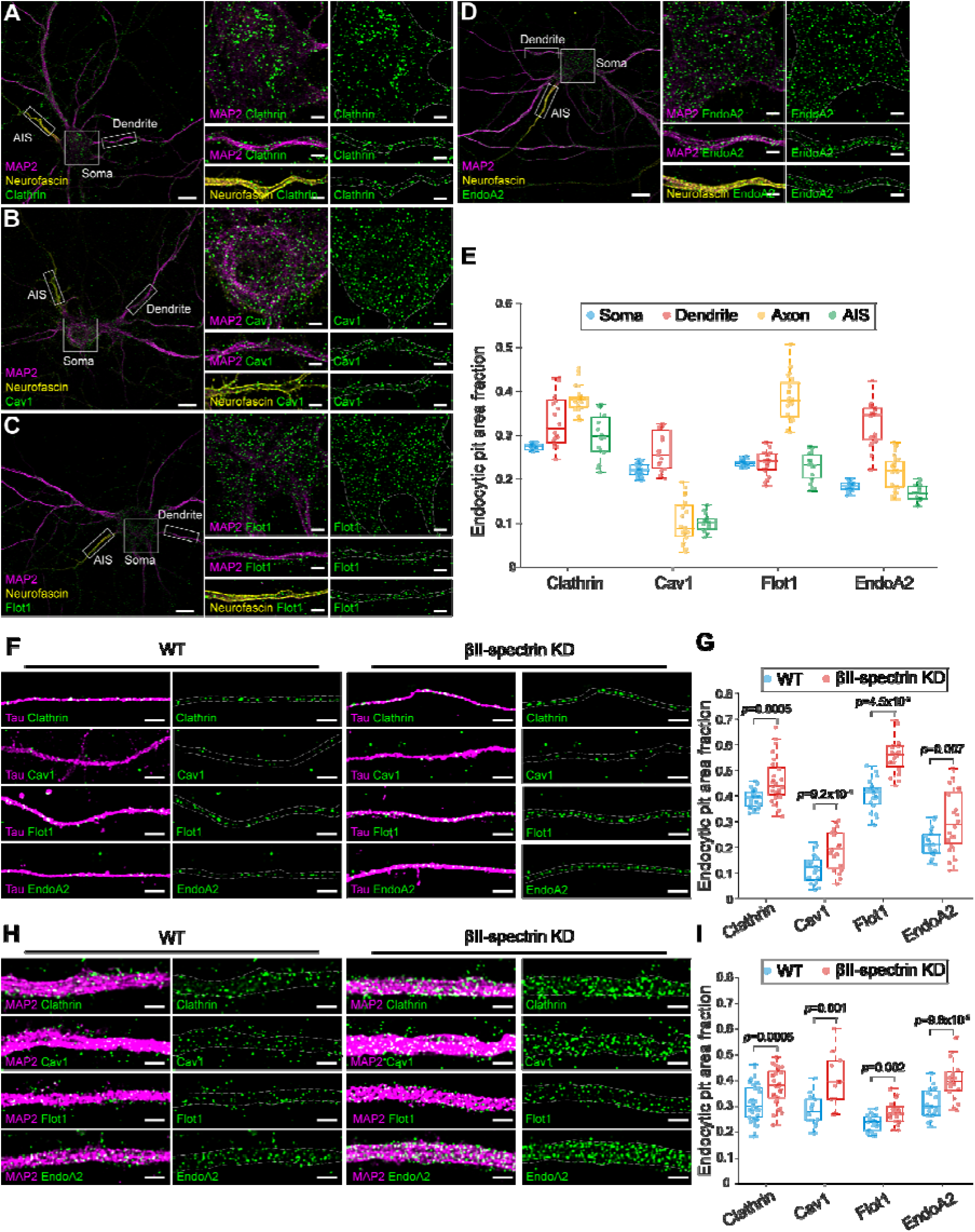
Compartment-resolved mapping of four endocytic pit types reveals broad suppression of basal endocytosis by the MPS in neurons. (**A-D**) Left: Stitched SIM images showing the distributions of endogenous endocytic pits, clathrin (**A**), Cav1(**B**), Flot1 (**C**) or EndoA2 (**D**) in WT neurons. Endocytic pits are shown in green, with compartment markers MAP2 (magenta) and neurofacsin (yellow). Scale bar: 10 µm. Right: Enlarged SIM images of the three boxed regions on the left, corresponding to soma, dendrite and AIS compartments, respectively. Scale bar: 2 µm. (**E**) Box plots showing the area fraction of endogenous endocytic pits in different compartments of WT neurons. (**F**) Left: SIM images of tau (magenta) and endogenous endocytic pits (green) in distal axons of WT neurons. Right: Same as Left, but in βII-spectrin knockdown (KD) neurons. Scale bars: 2 µm. (**G**) Box plots showing the area fraction of endogenous endocytic pits in distal axons of WT and βII-spectrin KD neurons. (**H**) Left: SIM images of MAP2 (magenta) and endogenous endocytic pits in dendrites of WT neurons. Right: Same as Left, but in βII-spectrin KD neurons. Scale bars: 2 µm. (**I**) Box plots showing the area fraction of endogenous endocytic pits in dendrites of WT and βII-spectrin KD neurons. All the experiments were replicated independently three times with similar results. Boxplots show the median and boundaries (first and third quartile), and the whiskers denote 1.5 times the interquartile range of the box. p-values calculated with two-sided unpaired Student’s t-test.

Having established the basal-level distributions of these endocytic pits across different neuronal compartments, we next aimed to examine the regulatory role of the MPS in modulating these distributions. A decade ago, super-resolution imaging enabled the discovery of the neuronal MPS, a structure in which actin filaments are organized into “rings” connected by spectrin tetramers, forming a one-dimensional (1D) periodic lattice with ∼190-nm spacing beneath the plasma membrane of neurites. In mature neurons, the MPS extends throughout ∼90% of axonal regions, including both the axon initial segment and distal axons, and is also observed in ∼50% of dendritic regions. The MPS is broadly distributed across diverse neuronal types, including excitatory and inhibitory neurons in both the central and peripheral nervous systems, and is conserved across species ranging from *C. elegans* to humans. This submembrane cytoskeletal network plays a critical role in organizing transmembrane proteins, such as ion channels, adhesion molecules, membrane transporters and receptors. The MPS has been implicated in enhancing axonal stability under mechanical stress, mediating mechanosensation, controlling neurite diameter and bundling(*31*), influencing axon degeneration, and regulating neuronal receptor signaling(*27*) and CME, emphasizing its essential role in neuronal function and health.

To study how MPS disruption affects the basal-level distributions and densities of the four types of endocytic pits, we transduced neurons with adenovirus expressing short hairpin RNA (shRNA) against βII-spectrin, a key structural component of the MPS, to disrupt the MPS both in the axonal and somatodendritic compartments, as described previously. The βII-spectrin expression level in βII-spectrin knockdown (KD) neurons decreased by ∼70%, compared to wild-type (WT) neurons, which were transduced with adenovirus expressing scrambled control shRNA as a control (**Fig. S2A, B**). The immunostained patterns for endogenous CCPs, Cav1-pits, Flot1-pits, and EndoA2-pits in WT and βII-spectrin KD neurons showed that MPS disruption significantly increased the densities of all four types of endocytic pits in all neuronal compartments examined including AIS, distal axonal segments, and dendrites, compared to WT neurons (**Fig. S2G, H and Fig. 1F, G, H, I**). Together, our results suggest that the MPS functions as a physical barrier to suppress basal endocytosis across all four endocytic pathways in all neuronal compartments, likely by acting as a physical barrier that restricts endocytic pit formation.

### Four major types of endocytic pits are localized within “clearings” of the MPS lattice in axons

We further investigated the structural basis underlying the MPS’s inhibitory role in regulating four major types of endocytosis in axons. To resolve the spatial relationship between endocytic pits and the MPS, we employed dual-color three-dimensional (3D) stochastic optical reconstruction microscopy (STORM), a super-resolution imaging technique with lateral resolution of 20–30 nm and axial resolution of 50–60 nm (**Fig. 2A**). The MPS was visualized in the cultured neurons (at or after DIV 21) through immunostaining of the C terminus of βII-spectrin or adducin, which mark the periodic “rings” formed by spectrin tetramer centers or actin filaments, respectively. Endocytic pits were immunostained using either antibodies against endogenous endocytic proteins, or an anti-GFP antibody for labeling the exogenously expressed GFP-tagged endocytic proteins through low-titer lentiviral transduction.

**Fig. 2.**
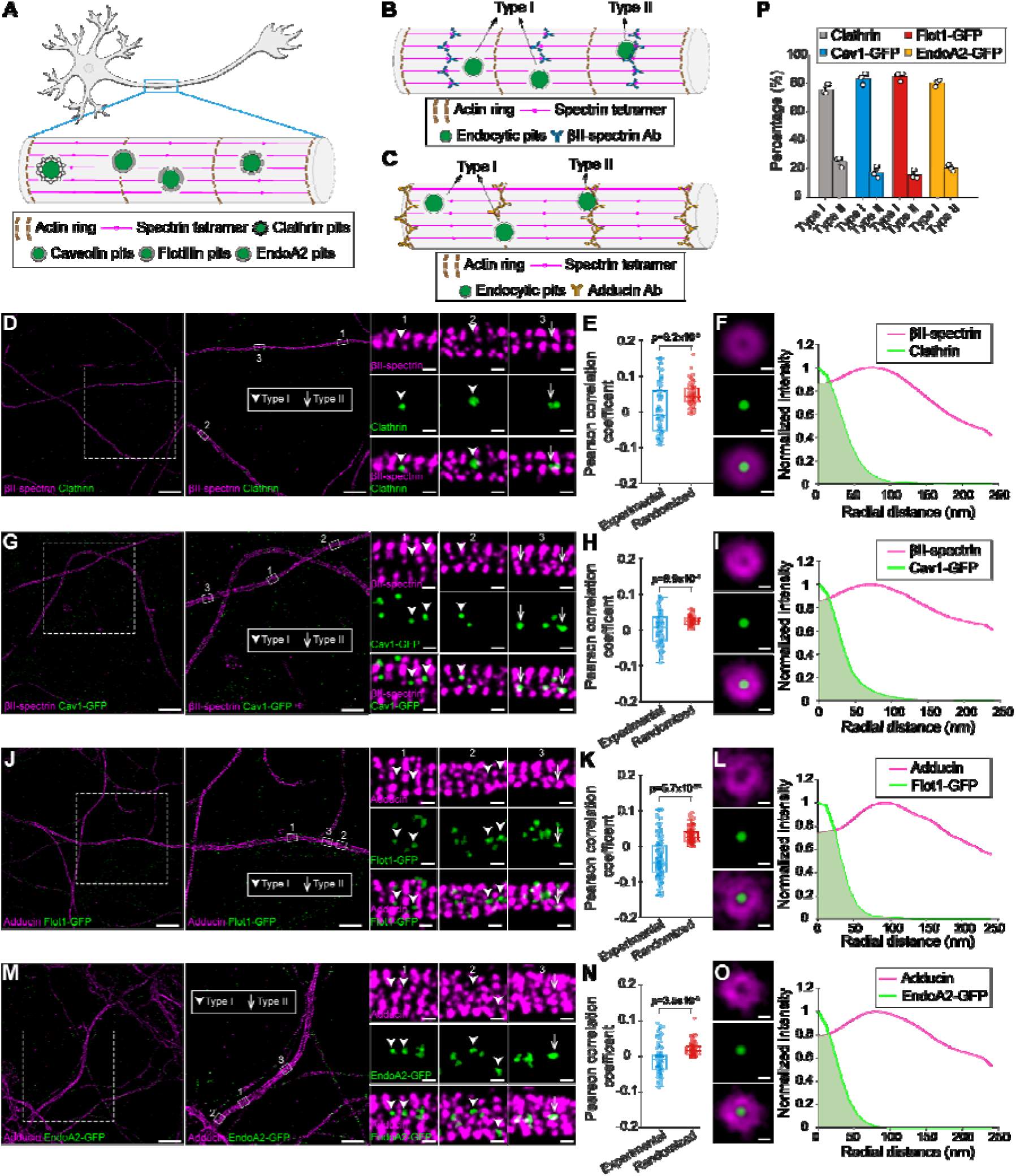
Four major types of endocytic pits are localized within “clearings” of the MPS lattice in axons. (**A**) Schematic illustrating the spatial distributions of clathrin, Cav1, Flot1 and EndoA2 endocytic pits, relative to periodic MPS lattice in axons. (**B**) Schematic illustrating two distinct types of endocytic pits based on their spatial positioning relative to periodic βII-spectrin lattice in axons. Type I pits do not overlap with MPS lattice, whereas Type II do. (**C**) Same as in b but showing spatial relationships with th periodic adducin lattice in axons. (**D**) Left: Dual-color STORM images of βII-spectrin (magenta) and endogenous clathrin (green) in axons. Right: Representative enlarged images of Type I and Type II clathrin-coated pits (CCPs) in the boxed regions. Scale bars: 10 µm (left), 5 µm (middle), 200 nm (right). (**E**) Pearson correlation coefficients between βII-spectrin and endogenous clathrin under experimental and randomized conditions. (**F**) Left: Averaged dual-color STORM images of βII-spectrin (magenta) and endogenous clathrin (green), generated by aligning individual STORM images to the centers of CCPs. Right: Radial intensity profiles of the averaged images shown on the left. Scale bar: 100 nm. (**G-I**) Same as in (**D–F**), but for βII-spectrin (magenta) and exogenously expressed Cav1 (green). (**J-L**) Same as in (**D–F)**, but for adducin (magenta) and exogenously expressed Flot1 (green). (**M-O**) Same as in (**D–F)**, but for adducin (magenta) and exogenously expressed EndoA2 (green). (**P**) Percentages of Type I and Type II pits for endogenous clathrin, exogenously expressed Cav1, exogenously expressed Flot1 and exogenously expressed EndoA2 in axons. Data are presented as mean ± s.e.m. (n = 3 biological replicates per condition). Boxplots show the median and boundaries (first and third quartile), and the whiskers denote 1.5 times the interquartile range of the box. p-values calculated with two-sided paired Student’s t-test.

3D STORM cross-sections (y/z views) revealed that all four types of endocytic pits were present both at the plasma membrane and within axonal shafts (**Fig. S3A**), with the fractions of endogenous endocytic pits being localized at the cell surface being 78.9 ± 4.0%, 82.3 ± 1.8%, 67.2 ± 1.7% and 67.0 ± 6.9% for CCPs, Cav1-pits, Flot1-pits, and EndoA2-pits, respectively (**Fig. S3B**). The surface-localized fractions of GFP-tagged endocytic pits closely resembled those of their endogenous counterparts, except for GFP-tagged Cav1 pits, which displayed a slightly lower surface-localized fraction, suggesting that both labeling strategies yield comparable results. To further validate our quantification method, we treated neurons with dyngo-4a, a potent dynamin inhibitor that prevents CCP scission from the plasma membrane(*35*), which led to an expected increase in the fraction of surface-localized CCPs in both axonal and dendritic compartments, confirming the reliability of our measurements (**Fig. S3D**).

Next, we quantified the diameters of all four types of endogenous endocytic pits localized at the plasma membrane, which averaged 82.5 ± 2.8, 75.0 ± 4.3, 71.7 ± 2.8 and 67.0 ± 3.4 nm for CCPs, Cav1-pits, Flot1-pits, and EndoA2-pits, respectively (**Fig. S3C**). The average diameters for the internalized fractions of the four endogenous endocytic pits were similar to those of surface-localized pits, indicating that the majority of surface-localized puncta analyzed are likely fully assembled endocytic pits (**Fig. S3C, E**). In addition, GFP-tagged endocytic pits exhibited similar diameters to their endogenous counterparts, for both surface-localized and internalized fractions (**Fig. S3C**).

For the surface-localized fractions of these four types of endocytic pits, we examined their spatial distributions relative to the MPS in axons by classifying the relative positions of endocytic pits into two scenarios (**Fig. 2B, C**): Type I) The pit has no overlap with the MPS; and Type II) the pit overlaps with the MPS. Quantification revealed that Type I pits were much more prevalent than Type II pits for all four types of endocytic pits, suggesting that all four types of pits are spatially excluded from the MPS (**Fig. 2D, G, J, M, P**). To confirm this exclusion, we calculated the Pearson’s correlation coefficients between the surface-localized pits and the MPS across 70-130 axonal regions, and found that these experimental coefficient values were significantly lower than those calculated from randomization controls, where the positions of all pits in each analyzed axonal region were randomly shuffled before calculating the Pearson’s correlation coefficient (**Fig. 2E, H, K, N**). As negative controls, we labeled neurons with two widely used membrane markers wheat germ agglutinin (WGA) and cholera toxin B subunit (CTB), and imaged their spatial distributions relative to the MPS in axons using STORM, following the same procedure as for the endocytic pits. The calculated Pearson’s correlation coefficients between either membrane marker and the MPS were not significantly different from those calculated from randomized controls, supporting the observed spatial exclusion between pits and the MPS as a true biological pattern (**Fig. S4A, B, C, D**). In addition, we compared the quantification results obtained from endogenously labeled pit proteins with those obtained from exogenously expressed GFP-tagged pit proteins (**Fig. S3E, F, G, H, I, J, K)**. Both labeling strategies yielded comparable results. Finally, to better visualize the average spatial pattern of these exclusion zones, we generated averaged STORM images aligned to the centers of endocytic pits. For all four types of pits, the corresponding radial fluorescence intensity profiles revealed that these pits are located in regions devoid of MPS immunostaining (**Fig. 2F, I, L, O**). These circular “clearing” structures observed closely resembled those CCP-centered “clearings” previously observed at the AIS(*8*). Together, these findings demonstrate that CCP-centered clearings are not only present at or near the AIS but also extend into distal axonal segments, and similar clearings are also present in the axons for the three clathrin-independent endocytic pits examined.

### Four major types of endocytic pits are localized within “clearings” of the MPS lattice in dendrites

The initial study reported that CCP-centered “clearings” observed at the AIS were absent from somatodendritic compartments. However, this absence may be attributed to imaging being performed on DIV14 neurons, while the MPS develops much more slowly in somatodendrites than in axons(*9*). The MPS coverage across the plasma membrane of somatodendrites reaches only ∼50% saturation around DIV 21–28, whereas MPS coverage in axons reaches ∼90% by DIV 14. This developmental delay raises the possibility that similar clearings may emerge in somatodendrites of more mature neurons (DIV 21 or older).

Since the MPS coverage is comparable between the soma and dendrites in mature neurons, and STORM imaging is technically easier in dendrites, we selected dendrites as the representative somatodendritic compartment for our imaging analysis (**Fig. 3A**). We first examined the spatial relationship between each of the four endocytic pit types and the surface-localized MPS in the radial direction within cross-section images of dendrites. The dendritic MPS was labeled by immunostaining either adducin or the N-terminus of βIII-spectrin (a β-spectrin isoform present primarily in somatodendrites but not axons), both of which mark the periodic “rings” formed by actin filaments (**Fig. 3B)**. Endocytic pits were immunostained using the same two labeling strategies employed for axon imaging. 3D STORM cross-sections (y/z views) revealed that all four types of endocytic pits were present both at the plasma membrane and within dendritic shafts (**Fig. S3F**), with the surface-localized fractions being similar to those observed in axons (**Fig. S3G**). The average diameters of the internalized endocytic pits closely matched those of the surface-localized fractions, and these average diameters determined in dendrites were comparable to those observed in axons. Both labeling strategies for endocytic pits produced consistent results (**Fig. S3H**).

**Fig. 3.**
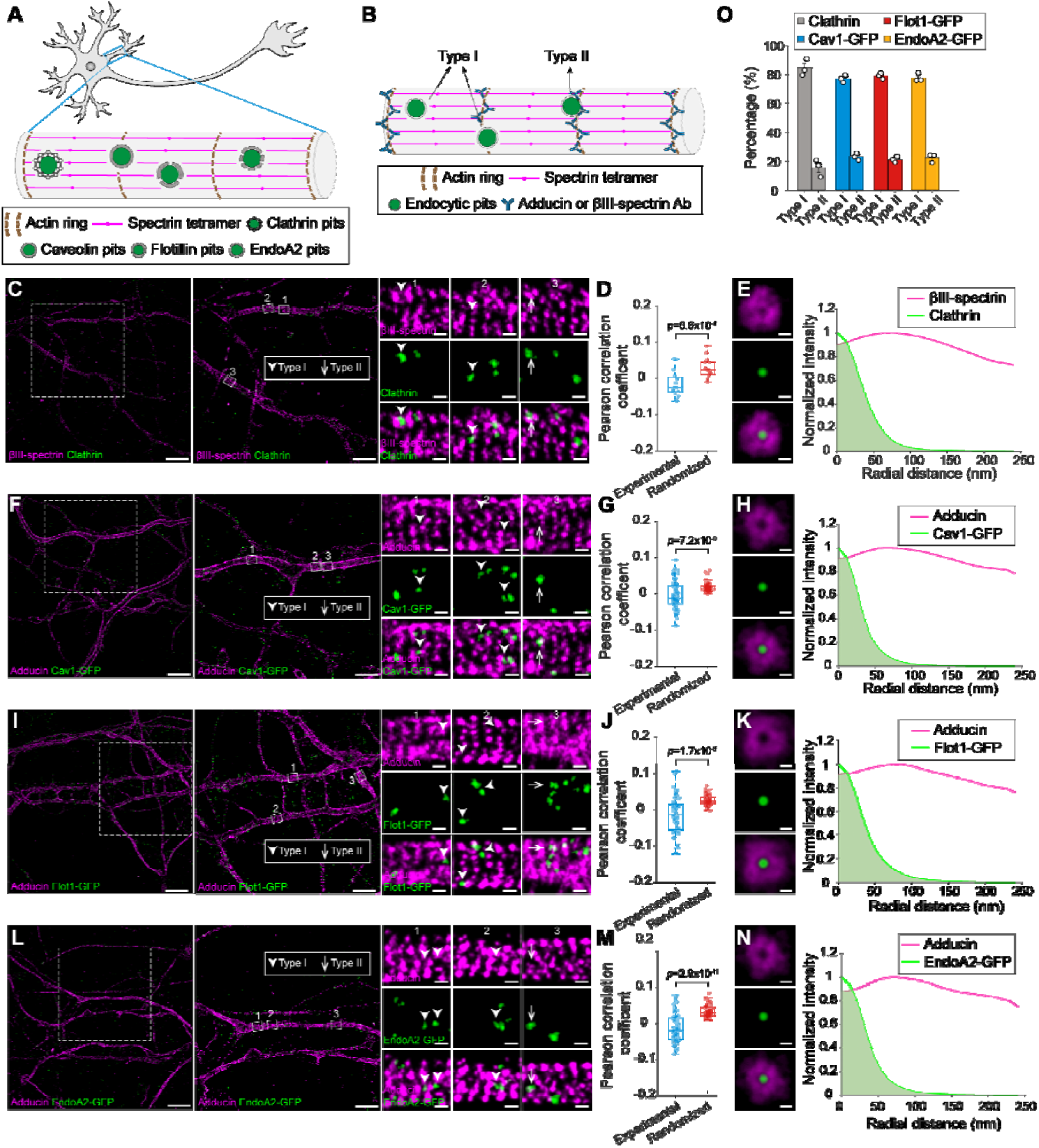
Four major types of endocytic pits are localized within “clearings” of the MPS lattice in dendrites. **(A**) Schematic illustrating the spatial distribution of clathrin, Cav1, Flot1 and EndoA2 endocytic pits, relative to periodic MPS lattice in dendrites. **(B**) Schematic illustrating two distinct type of endocytic pits based on their spatial positioning relative to periodic βIII-spectrin or adducin lattice in dendrites. Type I pits do not overlap with MPS lattice, whereas Type II pits do. **(C**) Left: Dual-color STORM images of βIII-spectrin (magenta) and endogenous clathrin (green) in dendrites. Right: Representative enlarged images of Type I and Type II clathrin-coated pits (CCPs) in the boxed regions. Scale bars: 10 µm (left), 5 µm (middle), 200 nm (right). **(D**) Pearson correlation coefficients between βIII-spectrin and endogenous clathrin under experimental and randomized conditions. (**E**) Left: Averaged dual-color STROM images of βIII-spectrin (magenta) and endogenous clathrin (green), generated by aligning individual STORM images to the centers of CCPs. Right: radial intensity profile of averaged images shown on the left. Scale bar: 100 nm. **(F-H**) Same as in (**C–E)**, but for adducin (magenta) and exogenously expressed Cav1 (green). **(I-K)** Same as in (**C–E**), but for adducin (magenta) and exogenously expressed Flot1 (green). (**L-N**) Same as in (**C–E**), but for adducin (magenta) and exogenously expressed EndoA2 (green). (**O**) Percentages of Type I and Type II endocytic pits for endogenous clathrin, exogenously expressed Cav1, exogenously expressed Flot1 and exogenously expressed EndoA2 in dendrites. Data is presented as mean ± s.e.m. (n = 3 biological replicates per condition). Boxplots show the median and boundaries (first and third quartile), and the whiskers denote 1.5 times the interquartile range of the box. p-values calculated with two-sided paired Student’s t-test.

We then examined the spatial relationship between each type of endocytic pit and the MPS within top-view STORM images of dendrites. As in the axon analysis, endocytic pits were classified into two types based on their spatial overlap with the MPS. Two-color STORM revealed that Type I pits (i.e., pits without overlap with the MPS) were markedly more prevalent than Type II pits for all four types of endocytic pits (**Fig. 3C, F, I, L, O and Fig. S5A, C, E, G**). The spatial exclusion pattern of endocytic pits from the MPS was confirmed by Pearson’s correlation analysis as performed for axons, where the experimental coefficients calculated from experimental images were significantly lower than those calculated from the corresponding randomization controls (**Fig. 3D, G, J, M and Fig. S5B, D, F**). Finally, intensity profiles from averaged STORM images aligned to the centers of endocytic pits revealed circular “clearings” surrounding clathrin-, Cav1-, Flot1- or EndoA2-endocytic pits in dendritic compartments (**Fig. 3E, H, K, N**). Together, these results suggest that circular “clearings” are a conserved structure in both axonal and somatodendritic compartments of mature neurons, with both clathrin-dependent and clathrin-independent endocytic pits residing at their centers.

### MPS inhibits ligand-induced endocytosis in both axonal and somatodendritic compartments

Given our observation that MPS disruption significantly increased the basal endocytosis of the four major types across all neuronal compartments, we investigated whether the MPS similarly inhibits ligand-induced endocytosis mediated by clathrin, caveolin, flotillin, or endophilin – which are generally considered to occur more rapidly than basal-level, constitutive endocytosis.

We first examined transferrin and low-density lipoprotein (LDL) endocytosis in neurons, both of which are forms of ligand-induced endocytosis primarily mediated by clathrin (i.e., CME) in most cell types. Incubating fluorophore-conjugated transferrin or LDL with cultured neurons resulted in an increased number of internalized transferrin or LDL over time in the somatodendrites but not axons, consistent with previous studies. To ensure that only internalized ligands were quantified, we performed an acid wash to remove surface-bound transferrin or LDL before imaging. To further validate the endocytic pathways involved in transferrin or LDL uptake, we pretreated neurons with four inhibitors targeting distinct pathways(*1*) and assessed their effects, including: 1) shRNA against clathrin heavy chain (CHC) to disrupt CME(*38*) with CHC knockdown confirmed by more than a 60% reduction in CHC immunofluorescence intensity in CHC KD neurons compared to WT neurons (**Fig. S6A, B**); 2) dyngo-4a, which inhibits CME, FEME and certain types of LRME; 3) nystatin, which disrupts lipid rafts and hence LRME by sequestering cholesterol; and 4) 5-[N-ethyl-N-isopropyl] amiloride (EIPA), a Na^+^/H^+^ exchanger inhibitor that disrupts macropinocytosis but not receptor-mediated endocytosis. Transferrin uptake by neurons was inhibited by CHC knockdown, dyngo-4a and EIPA but not by nystatin (**Fig. S6C, D**), suggesting that transferrin uptake depends on CME and possibly macropinocytosis. Consistent with this, imaging revealed that internalized transferrin strongly colocalized with clathrin but not with Cav1, Flot1, or EndoA2 (**Fig. S6E**), further supporting CME as the primary pathway of transferrin uptake. In contrast, LDL uptake was inhibited only by CHC knockdown and dyngo-4a, indicating that LDL uptake relies exclusively on CME (**Fig. S6H, I**). To determine whether MPS disruption affects clathrin-mediated transferrin and LDL uptake, we quantified the area fraction of internalized transferrin or LDL puncta, defined as the ratio of total transferrin or LDL puncta area to the corresponding dendritic area, in both WT and βII-spectrin KD neurons (**Fig. 4A and Fig. S6J**). By fitting the puncta area fraction over time to a single-exponential curve, we determined the time constants for uptake. Such time-course analysis revealed that transferrin uptake was faster in βII-spectrin KD neurons with a time constant τ=8.29 ± 0.37 min, compared to τ=17.15 ± 0.89 min in WT neurons (**Fig. 4B)**. Similarly, LDL uptake was enhanced in βII-spectrin KD neurons (τ = 35.25 ± 1.49 min), compared to WT neurons (τ = 54.63 ± 2.45 min) (**Fig. S6K)**. Notably, βII-spectrin knockdown did not alter surface expression levels of transferrin receptors or LDL receptors, ruling out increased surface expression levels of transferrin receptors or LDL receptors as a cause for the accelerated transferrin or LDL-induced endocytosis (**Fig. S6F, G, L, M**). Together, these findings indicate that the MPS acts as a physical barrier to restrict ligand-induced CME.

**Fig. 4.**
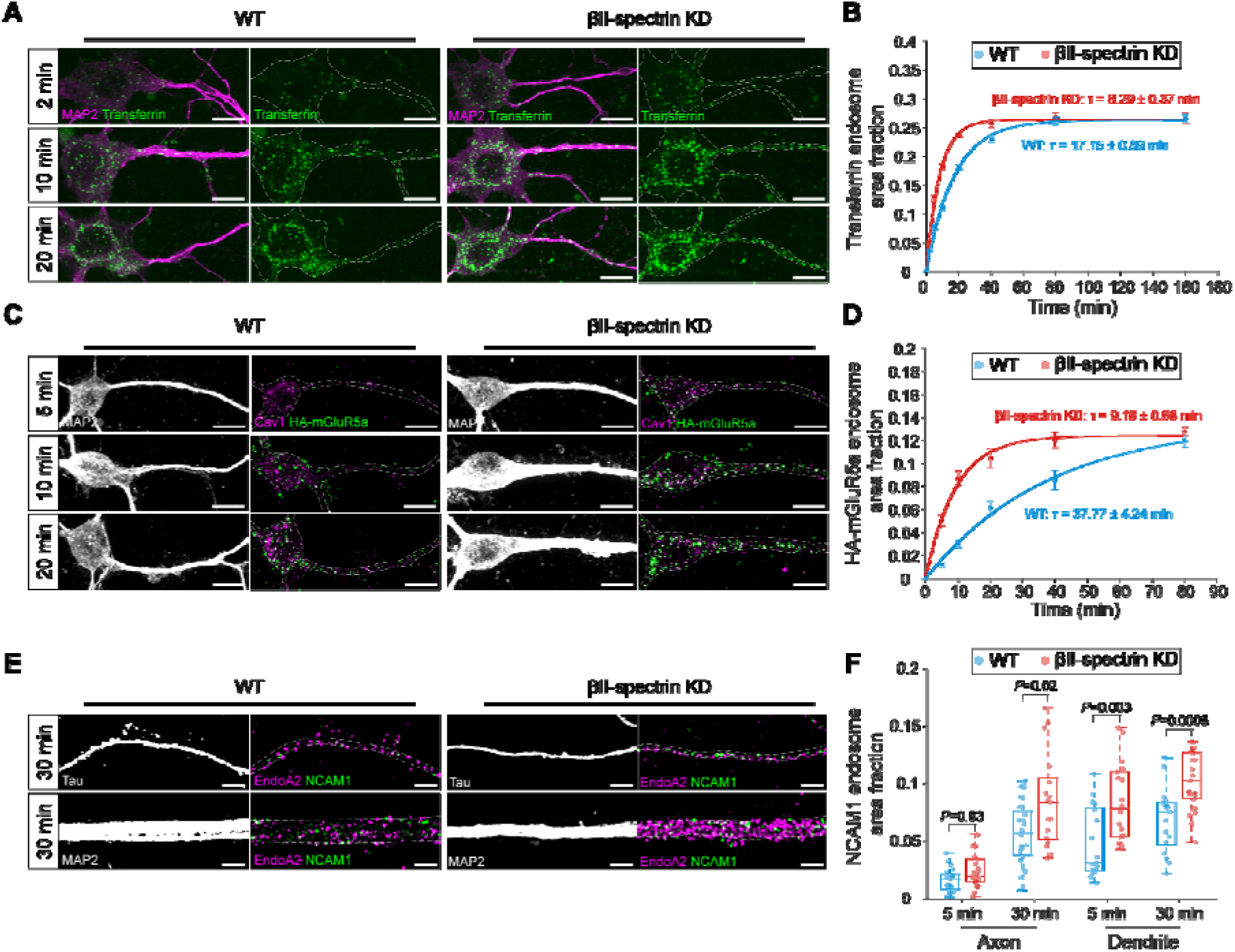
MPS inhibits ligand-induced endocytosis in both axonal and somatodendritic compartments. (**A**) Left: Confocal fluorescence images of MAP2 (magenta) and internalized CF568-transferrin (green) in somatodendritic region of WT neurons treated with CF568-transferrin for 2, 10, and 20 minutes. Right: Same as Left, but in βII-spectrin KD neurons. Scale bars: 10 µm. (**B**) Time course of CF568-transferrin endocytosis in somatodendritic regions of WT and βII-spectrin KD neurons, quantified by the area fraction of transferrin-positive endosomes. Solid lines represent single-exponential fits to the data. Data is presented as mean ± s.e.m. (**C**) Left: Confocal fluorescence images of MAP2 (gray), internalized HA-mGluR5a (green) and endogenous Cav1 (magenta) in somatodendritic region of WT neurons overexpressing HA-mGluR5a and treated with anti-HA antibody for 5, 10, and 20 minutes. Right: Same as Left, but in βII-spectrin KD neurons. Scale bars: 10 µm. (**D**) Time course of Cav1-mediated HA-mGluR5a endocytosis in the somatodendritic regions of WT and βII-spectrin KD neurons, quantified by the area fraction of Cav1-positive HA-mGluR5a endosomes. Solid lines represent single-exponential fit to the data. Data is presented as mean ± s.e.m. (**E**) Left: SIM images of internalized NCAM1 (green) and endogenous endoA2 (magenta) in axonal (top) and somatodendritic region (bottom) of WT neurons treated with anti-NCAM1 antibody for 30 minutes. Right: Same as Left, but in βII-spectrin KD neurons. Scale bars: 2 µm. (**F**) Boxplots of EndoA2-mediated NCAM1 endocytosis in axonal and somatodendriti regions of WT and βII-spectrin KD neurons, quantified by the area proportion of EndoA2-positive NCAM1 endosomes. Boxplots show the median and boundaries (first and third quartile), and the whiskers denote 1.5 times the interquartile range of the box. p-values calculated with two-sided unpaired Student’s t-test.

We next examined whether MPS disruption affects ligand-induced LRME. A previous study reported that in cultured neurons overexpressing HA-tagged metabotropic glutamate receptor 5a with the HA epitope tag located on its extracellular N-terminus (HA-mGluR5a), treatment with an anti-HA antibody induced HA-mGluR5a endocytosis across all neuronal compartments via a proposed lipid raft-mediated and clathrin-independent pathway(*39*). Using the same assay, treatment with the anti-HA antibody followed by an acid wash to remove surface-bound antibody resulted in a punctate immunostaining pattern of HA-mGluR5a, indicating its internalization. A subset of these puncta was colocalized with Cav1 pits but not with clathrin-, Flot1-, or EndoA2- pits (**Fig. S7A**), supporting a caveolin-mediated mechanism. To confirm the endocytic pathway involved, we tested the effects of the four inhibitors described above. Antibody-induced HA-mGluR5a endocytosis was significantly inhibited by dyngo-4a or nystatin, but not by CHC knockdown or EIPA, further indicating that HA-mGluR5a endocytosis occurs, at least partially, through caveolin-mediated endocytosis, a major LRME pathway (**Fig. S7B, C**). Next, we assessed the effect of MPS disruption on antibody-induced HA-mGluR5a endocytosis. WT and βII-spectrin KD neurons were incubated with the anti-HA antibody over a time course, followed by immunostaining for internalized HA-mGluR5a and endogenous Cav1. This allowed us to selectively analyze the fraction of internalized HA-mGluR5a that colocalized with endogenous Cav1-pitsa (**Fig. 4C**). Time-course analysis revealed that caveolin-mediated HA-mGluR5a endocytosis occurred faster in βII-spectrin KD neurons (τ=9.18 ± 0.68 min) compared to WT neurons (τ = 37.77 ± 4.24 min) (**Fig. 4D**). This enhancement was not due to changes in the surface expression of HA-mGluR5a (**Fig. S7D, E**). Together, our data indicates that MPS disruption facilitates ligand-induced LRME.

Finally, we assessed the impact of MPS disruption on ligand-induced FEME. When examining endogenous neural cell adhesion molecule 1 (NCAM1) internalization in mature neurons, triggered by treatment with a monoclonal anti-NCAM1 antibody against its extracellular domain, we found that the internalized NCAM1 puncta colocalized with EndoA2-pits but not with clathrin-, Cav1-, or Flot1-pits (**Fig. S7F**). Inhibitor experiments revealed that antibody-induced NCAM1 endocytosis was inhibited only by dyngo-4a, with no effect observed from CHC knockdown, nystatin, or EIPA (**Fig. S7G, H**). These results indicate that NCAM1 endocytosis occurs, at least partially, through endophilin-mediated endocytosis (i.e., FEME) in neurons. To assess the effect of MPS disruption, we compared the time course of NCAM1 endocytosis, specifically only for the fraction of NCAM1 puncta colocalized with endogenous EndoA2-pits, in WT and βII-spectrin KD neurons (**Fig. 4E**). Since βII-spectrin KD neurons exhibited ∼10% higher NCAM1 surface expression (**Fig. S7I, J)**, we selected neurite regions from both groups with comparable NCAM1 surface levels for our time course analysis. Our results showed that the internalization of the NCAM1 pool that is mediated by EndoA2 was increased in both axonal and dendritic compartments of βII-spectrin KD neurons compared to WT neurons (**Fig. 4F**), demonstrating that the MPS also restricts ligand-induced FEME. Together, our findings demonstrate that the MPS serves as a regulatory barrier that inhibits both clathrin-dependent and clathrin-independent ligand-induced endocytosis in axonal and somatodendritic compartments.

### The MPS acts as a dynamic barrier modulated by endocytosis and ERK signaling through a positive feedback mechanism

We next investigated whether ligand-induced endocytosis can, in turn, affect the MPS. Previously, we demonstrated that ligand binding to the extracellular domain of cannabinoid type 1 receptor (CB1) or NCAM1 induces the activation of the extracellular signal–regulated kinase (ERK), a key cell signaling pathway that involved in neuronal development and stress response, and hippocampal learning and survival(*40*). While we observed that CB1 ligand-induced ERK activation resulted in calpain-dependent MPS degradation, it remains unclear whether this phenomenon is common across other ligand-receptor systems (**Fig. 5A**). Using an IF-based ERK activation assay previously developed to monitor CB1- and NCAM1-induced ERK activation, we quantitatively assessed phosphorylated ERK (pERK) levels at different time points after ligand stimulation of three receptors: transferrin receptor (TfR), HA-tagged mGluR5a, and NCAM1. All ligand treatments induced ERK activation, which persisted for up to 60 minutes post-treatment (**Fig. 5B, C**). Inhibiting ligand-induced TfR, HA-mGluR5a and NCAM1endocytosis using dyngo-4a significantly reduced ERK activation (**Fig. 5B, C)**, indicating that a substantial portion of ERK activation occurs at internalized endosomes, and that ligand-induced endocytosis of TfR, mGluR5a and NCAM1 effectively contributes to ERK signaling. Notably, this contrasts with CB1 ligand-induced ERK activation, which does not rely on CB1 endocytosis and instead appears to occur primarily at the cell surface rather than in endosomes.

**Fig. 5.**
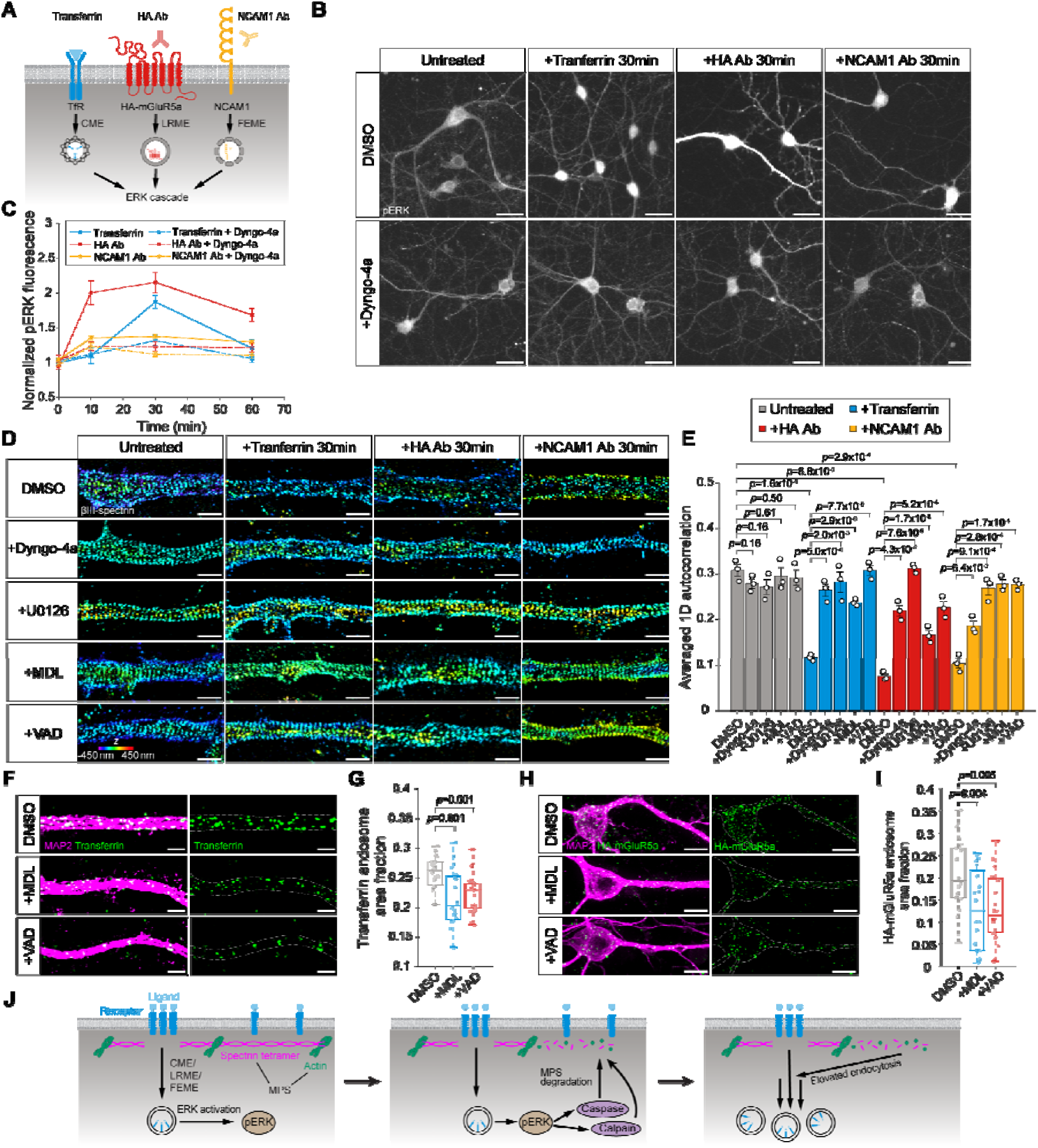
The MPS acts as a dynamic barrier modulated by endocytosis and ERK signaling through a positive feedback mechanism. (**A**) Schematic illustrating ligand-induced activation of ERK signaling via three major endocytic pathways: CME of transferrin receptor, LRME of HA-mGluR5a, and FEME of NCAM1 upon respective ligand binding. (**B**) Top: Epi-fluorescence images showing pERK immunostaining in neurons without ligand treatment, neurons treated with CF568-transferrin or anti-HA antibody treatmentfor 30 minutes, and neurons overexpressing HA-mGluR5a treated with anti-HA for 30 minutes. Bottom: Same as the top, but with neurons pretreated with dyngo-4a for 30 minutes before ligand treatment. Scale bars: 25 µm. (**C**) Time course of ERK activation in WT neurons treated with CF568-transferrin or anti-HA antibody, and in WT neurons overexpressing HA-mGluR5a treated with anti-HA antibody, with and without dyngo-4a pre-incubation. Data is presented as mean ± s.e.m. (**D**) 3D STORM images of immunostained βIII-spectrin in dendrites of neurons under various treatments. First column: neurons pretreated with DMSO (control), dyngo-4a, U0126 (a MEK/ERK inhibitor), MDL-28170 (MDL; a calpain inhibitor), or Z-VAD-FMK (VAD; a caspase inhibitor) alone. Second column: neurons pretreated with the same inhibitors followed by 30-minute treatment of CF568-transferrin. Third column: neurons overexpressing HA-mGluR5a pretreated with the same inhibitors followed by 30-minute treatment of anti-HA antibody. Fourth column: neurons pretreated with the same inhibitors followed by 30-minute treatment of anti-NCAM1 antibody. Scale bars: 1 µm. Color scale bar represents the z-coordinate information. (**E**) Averaged 1D autocorrelation amplitudes of βIII-spectrin in neurons, calculated for the same conditions as in (**D**). (**F**) SIM images of MAP2 (magenta) and internalized CF568-transferrin (green) in neurons pretreated with DMSO, MDL or VAD followed by 40-minute treatment of CF568-transferrin. Scale bars: 2 µm. (**G**) Boxplot of transferrin endocytosis in neurons, quantified by the area fraction of transferrin-positive endosomes. (**H**) Confocal fluorescence images of MAP2 (magenta) and internalized HA-mGluR5a (green) in neurons overexpressing HA-mGluR5a pretreated with DMSO, MDL or VAD followed by treatment of anti-HA antibody for 40 minutes. Scale bars: 10 µm. (**I**) Box plot of HA-mGluR5a endocytosis in neurons, quantified by the area fraction of HA-mGluR5a endosomes. (**J**) Schematic summarizing the proposed feedback mechanism: receptor endocytosis via CME, LRME, or FEME activates ERK signaling, which triggers calpain- and caspase-mediated MPS degradation; MPS disruption in turn facilitates further endocytosis, establishing a positive feedback loop. Boxplots show the median and boundaries (first and third quartile), and the whiskers denote 1.5 times the interquartile range of the box. p-values calculated with two-sided unpaired Student’s t-test.

We next investigated whether ligand-induced endocytosis and ERK activation leads to protease-dependent MPS degradation. Using average 1D autocorrelation amplitude analysis, a quantification method we previously developed to measure the integrity of MPS, we examined the effects of ligand-induced TfR, HA-mGluR5a, or NCAM1 endocytosis on MPS structure in neurons. Quantitative analysis of 3D STROM images of βIII-spectrin revealed significant MPS degradation following ligand stimulation, as evidenced by decreased average 1D autocorrelation amplitudes (**Fig. 5D, E**). Inhibiting ligand-induced TfR, HA-mGluR5a, or NCAM1 endocytosis using dyngo-4a prevented MPS degradation (**Fig. 5D, E)**. Preincubation with U0126, an inhibitor of MEK, the kinase upstream of ERK, also protected the MPS from degradation (**Fig. 5D, E)**. These results indicate that both endocytosis and ERK activation are upstream events required for MPS degradation. To identify the proteases involved, we tested the effects of MDL-28170 (a calpain inhibitor) and Z-VAD-FMK (a caspase inhibitor), both of which are known to cleave brain spectrin(*41*) and can be activated by ERK. Treatment of neurons with either protease inhibitor rescued MPS integrity, with varying degrees of protection, suggesting that endocytosis-induced MPS degradation is calpain- and caspase-dependent (**Fig. 5D, E**).

To further probe whether increased endocytosis alone could degrade the MPS, we investigated whether moderate overexpression of GFP-tagged endocytic pit proteins (i.e., Cav1, Flot1, and EndoA2) which may increase endocytic pit density, could similarly affect MPS integrity. We first confirmed that overexpressing these endocytic pit proteins in neurons significantly increased the endocytic pit area fractions compared to WT neurons, as quantified from the 3D STORM images described above (**Fig. S8A**). We then assessed MPS integrity using epi-fluorescence and STORM imaging. Compared to WT neurons and neurons overexpressing a plasma membrane-associated protein BSAP1 (an endocytic pit-irrelevant control), neurons overexpressing GFP-tagged Cav1, Flot1, or EndoA2 exhibited a moderate decrease in both the average βII-spectrin immunofluorescence intensity (quantified by epi-fluorescence imaging) and the average 1D autocorrelation amplitude (quantified by STORM imaging) in axons and dendrites (**Fig. S8B, C, D, E**). These findings suggest that increased endocytic pit density and enhanced endocytosis activity alone can induce moderate MPS degradation, further supporting our model that endocytosis contributes to MPS disruption.

So far, our data demonstrates that the MPS suppresses endocytosis and conversely, endocytosis can lead to MPS degradation. This reciprocal relationship suggests that endocytosis-induced MPS degradation may be a programmed cellular mechanism designed to rapidly remove the MPS barrier, facilitating accelerated endocytosis once triggered. To further test this hypothesis, we examined whether preventing MPS degradation could suppress ligand-induced endocytosis. For both ligand-induced TfR and HA-mGluR5a endocytosis, we quantified the internalized endocytic pit area fractions following ligand treatment, with or without calpain or caspase inhibitor treatment. Neurons were treated with either protease inhibitor, both prior to ligand addition and continuously throughout the endocytosis process. Compared to untreated conditions, inhibiting MPS degradation via calpain or caspase inhibitors significantly reduced ligand-induced TfR and HA-mGluR5a endocytosis (**Fig. 5F, G, H, I**). These results support our model that the MPS acts as a dynamic physical barrier to suppress endocytosis. When rapid endocytosis is required, cells activate ERK-protease pathways to induce calpain- and caspase-dependent MPS degradation, thereby removing the MPS barrier and promoting faster receptor internalization. This mechanism establishes a previously unknown positive feedback loop, wherein endocytosis drives MPS degradation, which in turn accelerates subsequent endocytosis and amplifies downstream signaling to support rapid neuronal responses (**Fig. 5J**).

### The MPS serves as a neuroprotective barrier by restricting APP endocytosis and A**β**42 accumulation

Increasing evidence indicates that dysregulated endocytosis in neurons contributes to the pathogenesis of Alzheimer’s disease (AD), a progressive neurodegenerative disorder where dementia symptoms gradually worsen over a number of years. Elevated production and accumulation of misfolded β-amyloid peptide (Aβ) in the brain correlates with neuronal aging and the AD development. The pathogenic form of Aβ (Aβ42) is generated from amyloid precursor protein (APP) after cell surface-localized APP undergoes endocytosis and is processed by sequential β- and γ-secretase cleavage in neuronal endosomes and lysosomes(*44*) (**Fig. 6A**). This raises the possibility that the MPS may have a neuroprotective role by suppressing APP endocytosis, thereby limiting Aβ production and accumulation.

**Fig. 6.**
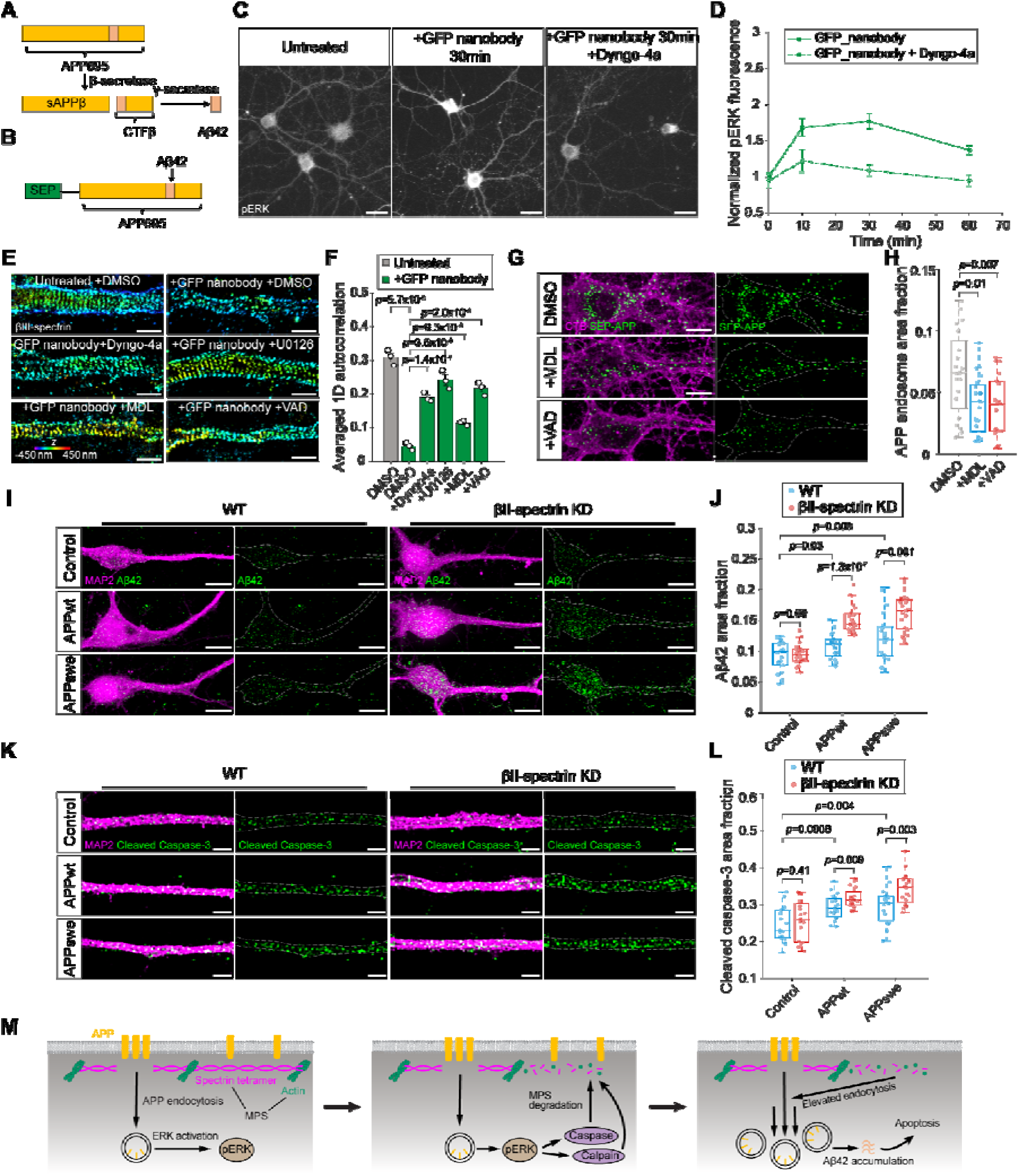
The MPS serves as a neuroprotective barrier by restricting APP endocytosis and Aβ42 accumulation. (**A**) Schematic illustrating Aβ42 production from APP695. APP695 is first cleaved by β-secretase, generating sAPPβ and CTFβ. CTFβ is further cleaved by γ-secretase to produce Aβ42. (**B**) Schematic illustrating the structure of SEP-APP, of which the N-terminal GFP variant SEP is fused to APP695. (**C**) Left: Epi-fluorescence images of pERK in neurons overexpressing SEP-APP without ligand treatment. Middle: Same as left, but treated with GFP nanobody for 30 minutes. Right: Same as middle, but with dyngo-4a pre-incubation before GFP nanobody treatment. Scale bars: 25 µm. (**D**) Time course of ERK activation in neurons overexpressing SEP-APP treated with GFP nanobody, with and without dyngo-4a pre-incubation. Data is presented as mean ± s.e.m. (**E**) 3D STORM images of immunostained βIII-spectrin in dendrites of neurons pretreated with DMSO, dyngo-4a, U0126, MDL-28170 (MDL) or Z-VAD-FMK (VAD) followed by treatment of GFP nanobody for 30 minutes. Scale bars: 1 µm. Color scale bar represents the z-coordinate information. (**F**) Averaged 1D autocorrelation amplitude of βIII-spectrin in neurons, calculated for the same conditions as in (**E**). (**G**) Confocal fluorescence images of CTB (magenta) and internalized SEP-APP (green) in neurons pretreated with DMSO, MDL or VAD followed by treatment of GFP nanobody for 40 minutes. Scale bars: 10 µm. (**H**) Boxplots of SEP-APP endocytosis in neurons, quantified by the area fraction of SEP-APP endosomes. (**I**) Left: Confocal fluorescence images of MAP2 (magenta) and intracellular Aβ42 (green) in WT neurons, neurons overexpressing APPwt, and neurons overexpressing APPswe. Right: Same as left, but in βII-spectrin KD neurons. Scale bars: 10 µm. (**J**) Boxplots of intracellular Aβ42 in somatodendritic regions of neurons, quantified by the area fraction of Aβ42. (**K**) Left: SIM images of MAP2 (magenta) and cleaved caspase-3 (green) in WT neurons, neurons overexpressing APPwt, and neurons overexpressing APPswe. Right: Same left, but in βII-spectrin KD neurons. Scale bars: 2 µm. (**L**) Boxplots of cleaved caspase-3 in dendrites of neurons, quantified by the area fraction of cleaved caspase-3. (**M**) Schematic illustrating APP endocytosis triggers downstream ERK signaling, leading to MPS degradation through caspase- and calpain-mediated spectrin cleavage. This degradation further accelerates APP endocytosis, promoting intracellular Aβ42 accumulation and activation of caspase-3, the marker for neuronal apoptosis. Boxplots show the median and boundaries (first and third quartile), and the whiskers denote 1.5 times the interquartile range of the box. p-values calculated with two-sided unpaired Student’s t-test.

To test this hypothesis, we employed a previously established ligand-induced APP endocytosis system(*45*) to manipulate and monitor APP endocytosis in neurons. In this system, a GFP variant, super-ecliptic pHluorin (SEP), was fused to the N-terminus of APP695, the predominant APP isoform in neuronal tissues (**Fig. 6B**). Treating neurons overexpressing SEP-APP with a fluorophore-conjugated nanobody (∼ 15 kDa) against SEP induced SEP-APP endocytosis in the somatodendritic compartments(*45*). Pretreating neurons with our four described inhibitors revealed that SEP-APP endocytosis involves complex pathways, proceeding through both clathrin-dependent and clathrin-independent pathways (**Fig. S9A, B**). This finding contrasts with a previous study that identified nanobody-induced SEP-APP endocytosis as strictly clathrin-independent(*45*), potentially due to differences in experimental conditions. We then compared SEP-APP endocytosis rates in both WT and βII-spectrin KD neurons. SEP-APP uptake was significantly faster in βII-spectrin KD neurons (τ = 14.52 ± 0.38 min) compared to WT neurons (τ = 37.21 ± 5.11 min), suggesting that MPS degradation accelerated APP internalization (**Fig. S9C, D**). Notably, this endocytosis enhancement was not due to an increased APP surface expression upon βII-spectrin knockdown (**Fig. S9E, F**). Similar to other ligand-induced endocytosis observed in this study, SEP-APP endocytosis activated sustained downstream ERK signaling, as indicated by increased pERK immunofluorescence (**Fig. 6C, D**). Following the nanobody-induced SEP-APP endocytosis, significant MPS degradation was observed, shown by reduced 1D autocorrelation amplitude of the periodic βIII-spectrin distribution (**Fig. 6E, F**). Inhibiting SEP-APP endocytosis using dyngo-4a or U0126 prevented MPS degradation, indicating the MPS degradation is endocytosis- and ERK-dependent. Moreover, this MPS degradation was partially prevented by calpain or caspase inhibitor treatment, indicating that MPS degradation was mediated by calpain- and caspase-dependent spectrin cleavage (**Fig. 6E, F**). Conversely, protecting MPS from degradation using calpain or caspase inhibitor significantly reduced SEP-APP endocytosis in neurons (**Fig. 6G, H**), reinforcing the role of the MPS as a regulatory barrier that restricts APP internalization. These findings suggested that APP endocytosis promotes ERK signaling, which facilitates MPS degradation, establishing a positive feedback loop that accelerates APP internalization.

We next examined whether increased APP endocytosis in MPS-disrupted neurons leads to elevated Aβ production, a hallmark of AD. To avoid potential confounding effects from SEP tagging, we quantified the Aβ accumulation in cultured mature neurons with and without overexpressing wild-type APP695 (APPwt) or APP695 bearing the K670M/N671L Swedish mutation (APPswe), a familial AD-associated variant known to enhance Aβ accumulation. We employed a previously developed IF-based Aβ42 quantification assay using a highly specific anti-Aβ42 C-terminal antibody(*46*). Immunostaining of intracellular Aβ42, the key pathogenic form of Aβ, revealed elevated Aβ42 levels in neurons overexpressing either APPwt or APPswe, compared to neurons without APP overexpression (i.e., WT neurons), confirming that overexpressing APPwt and APPswe lead to increased intracellular Aβ42 production(*47*) (**Fig. 6I, J)**. Interestingly, βII-spectrin KD neurons overexpressing APPwt or APPswe exhibited significantly higher levels of intracellular Aβ42 accumulation than WT neurons overexpressing APPwt or APPswe (**Fig. 6I, J)**. It is important to note that Aβ42 levels remained comparable between WT and βII-spectrin KD neurons without APP overexpression, indicating that the observed increase in Aβ42 accumulation requires both MPS disruption and APP overexpression. Together, these findings suggest that the MPS plays a neuroprotective role in limiting Aβ42 accumulation, likely by suppressing APP endocytosis and reducing subsequent secretase-mediated APP cleavage in endosomes and lysosomes of neurons.

Intraneuronal Aβ42 accumulation occurs before the formation of extracellular Aβ plaques and is known to trigger a cascade of pathological events, including proteasome and mitochondrial dysfunction, calcium dysregulation, and synaptic impairment(*48*). Since intraneuronal Aβ42 accumulation has been previously linked to activation of caspase-3, a canonical marker of apoptosis, as well as neuronal apoptosis, we investigated whether MPS disruption exacerbates Aβ42-mediated neurotoxicity by comparing caspase-3 activations levels in WT and βII-spectrin KD neurons with and without APPwt or APPswe overexpression. Immunostaining for cleaved (activated) caspase-3 using a previously reported, highly specific anti-caspase-3 antibody(*51*) revealed significantly higher caspase-3 activation in neurons overexpressing APPwt or APPswe compared to non-overexpressing controls, with even greater caspase-3 activation observed in βII-spectrin KD neurons (**Fig. 6K, L**), supporting the hypothesis that MPS disruption accelerates APP internalization and subsequent Aβ42 production, thereby promoting neurotoxicity and neuronal apoptosis. Importantly, the observed patterns of caspase-3 activation mirrored the corresponding Aβ42 accumulation levels across WT and βII-spectrin KD neurons with and without APP overexpression, suggesting that elevated apoptosis signals are likely a direct consequence of increased Aβ42 accumulation. Together, these findings support a model in which the MPS acts as a neuroprotective barrier that restricts APP endocytosis, thereby limiting intraneuronal Aβ42 production, accumulation and subsequent neuronal apoptosis (**Fig. 6M**). When the MPS degradation occurs, previously observed in both aged and the neurodegenerative conditions(*52*), APP endocytosis accelerates, leading to pathological Aβ42 accumulation, neuronal dysfunction, and ultimately neuronal death. This mechanism provides insight into how MPS degradation may contribute to the progression of Alzheimer’s disease, and highlights preserving MPS integrity as a potential therapeutic strategy to mitigate APP-related neurodegeneration.

## Discussion

In this study, we leveraged super-resolution fluorescence imaging to visualize the structural interplay between the MPS and endocytic events at nanoscale resolution, and identified endocytic “clearing” structures analogous to those previously reported for CCPs at the AIS and proximal axon(*8*). Notably, we extended these observations to distal axonal and somatodendritic compartments, which were not reported in the initial study(*8*). The absence of such structures in the initial study may be due to the use of immature neurons, in which the somatodendritic MPS had not yet developed into a sufficiently dense and continuous lattice network(*9*). Importantly, we found that these clearing structures are not unique to CME but also observed for other major endocytic pathways, including LRME, such as caveolin- and flotillin-mediated internalization, as well as FEME. These findings reveal that MPS-based clearings represent a conserved structural feature across multiple endocytic mechanisms. Our super-resolution imaging thus provides direct structural evidence that the MPS functions as a spatial gatekeeper of endocytosis, regulating access through discrete clearings embedded within the periodic actin-spectrin lattice.

Aligned with our structural findings, functional assays comparing the endocytosis rates in neurons with and without MPS disruption support the role of the MPS as a universal regulator of all major endocytic pathways, including CME, LRME, and FEME. Disruption of the MPS not only resulted in a pronounced elevation of the intrinsic, basal-level endocytic activity across multiple pathways, but also markedly accelerated ligand-induced internalization events mediated through CME, LRME, and FEME, underscoring the MPS’s central role as a structural suppressor of both constitutive and stimulus-evoked endocytic processes. Importantly, our data reveals that ligand-induced endocytosis itself activates a positive feedback loop that further accelerates endocytic uptake: Ligand-induced endocytosis triggers sustained ERK activation, which in turn activates calpain and caspase to cleave spectrin and degrade the MPS; This degradation process effectively removes the MPS-imposed barrier, thereby facilitating faster and more robust endocytic uptake. The presence of this positive feedback loop illustrates a finely tuned mechanism whereby neurons can rapidly enhance endocytosis when rapid signaling or cargo internalization is required. This dynamic interplay underscores the MPS as both a gatekeeper and a responsive element in endocytic regulation.

In addition to CME, our findings reveal that other major endocytic pathways, including LRME and FEME, also occur at the AIS, distal axon, and somatodendritic compartments of neurons under basal (unstimulated) conditions. This broad spatial distribution underscores the pervasive and compartmentally integrated nature of endocytosis in mature neurons, extending beyond the traditionally emphasized role of CME. By using a panel of well-characterized ligands and assessing their colocalization with pathway-specific endocytic structures, we were able to selectively probe the activity of distinct endocytic mechanisms: transferrin and LDL for CME, mGluR5a for LRME, and NCAM1 for FEME. These assays provide a robust framework for the investigation of specific endocytic pathways in neurons.

Our findings also reveal important implications for neuronal health and disease. We observed that MPS degradation accelerates APP endocytosis, leading to increased production and intracellular accumulation of Aβ42, a pathogenic hallmark of Alzheimer’s disease. The elevated intraneural Aβ42 levels resulting from increased APP internalization were accompanied by downstream neurotoxic events, such as increased activation of caspase-3, which has been observed in Alzheimer’s brain(*53*). Stabilizing the MPS using calpain or caspase inhibitors reduced APP endocytosis, suggesting that the MPS serves as a neuroprotective structure that restricts excessive APP internalization and potentially slows down subsequent Aβ42 generation and caspase-3 activation. Moreover, the positive feedback loop we identified—whereby endocytosis triggers ERK activation, leading to protease-mediated MPS degradation—provides a plausible mechanism through which neurodegeneration may be exacerbated. Supporting this idea, spectrin cleavage products have been proposed as biomarkers for aging and neurodegenerative diseases(*54*). In aging neurons or those subjected to chronic stress, MPS instability may initiate a self-perpetuating cycle of heightened endocytosis and sustained ERK signaling, further accelerating APP internalization and Aβ42 accumulation. This insight is consistent with growing evidence that dysregulated endocytosis and protease activation contribute to Alzheimer’s pathogenesis.

Together, these findings position the MPS as a central structural regulator of neuronal endocytosis, capable of both constraining and dynamically modulating membrane trafficking across diverse cellular contexts. By uncovering its roles in both physiological regulation and disease-relevant pathways, our study not only expands the conceptual framework of endocytic control in neurons but also highlights the MPS as a potential therapeutic target for modulating endocytosis in neurodegenerative disorders.

## Author contributions

R.Z. conceived the project. J.F. and R.Z. designed experiments, interpreted the data and wrote the manuscript. J.F., Y.Z, C.L. performed experiments. J.F. analyzed the data. Y.T. provided coding assistance during the early stage of the project. R.Z. supervised the project.

## Data availability

All data supporting the findings of this study are provided within the paper and its supplemental information.

## Code availability

Custom Python and MATLAB codes for image acquisition and STORM analysis are available at https://github.com/ZhuangLab (DOI: 10.5281/zenodo.3264857)(*61*). Other custom MATLAB codes used in this study are available from the corresponding authors upon reasonable request.

## Supplementary Materials for

**Fig. S1.**
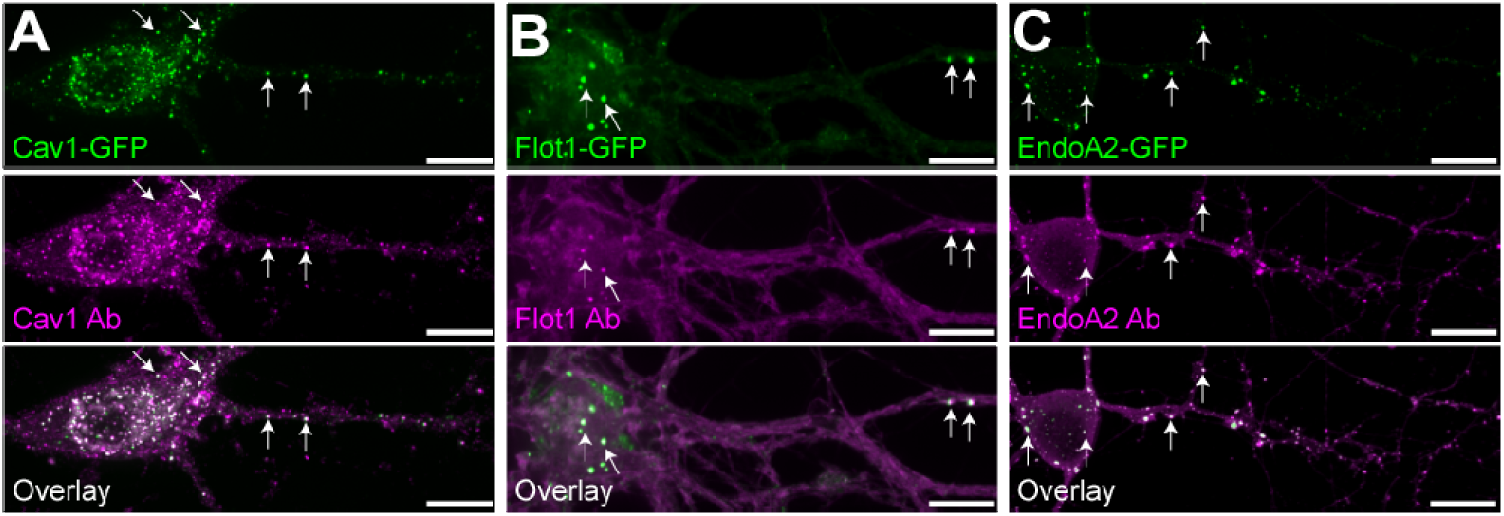
Validation of antibodies and immunofluorescence-based labeling of caveolin-, flotillin-, and endophilinA2-positive endocytic pits, related to Fig. 1. (**A**) Confocal fluorescence images of neurons exogenously expressing GFP-tagged Cav1 and immunostained for Cav1. Regions of strong colocalization are indicated by white arrows. (**B**) Same as (**A**), but for neurons exogenously overexpressing GFP-tagged Flot1 and immunostained for Flot1. (**C**) Same as (**A**), but for neurons exogenously expressing GFP-tagged EndoA2 and immunostained for EndoA2. Scale bars: 10 µm.

**Fig. S2.**
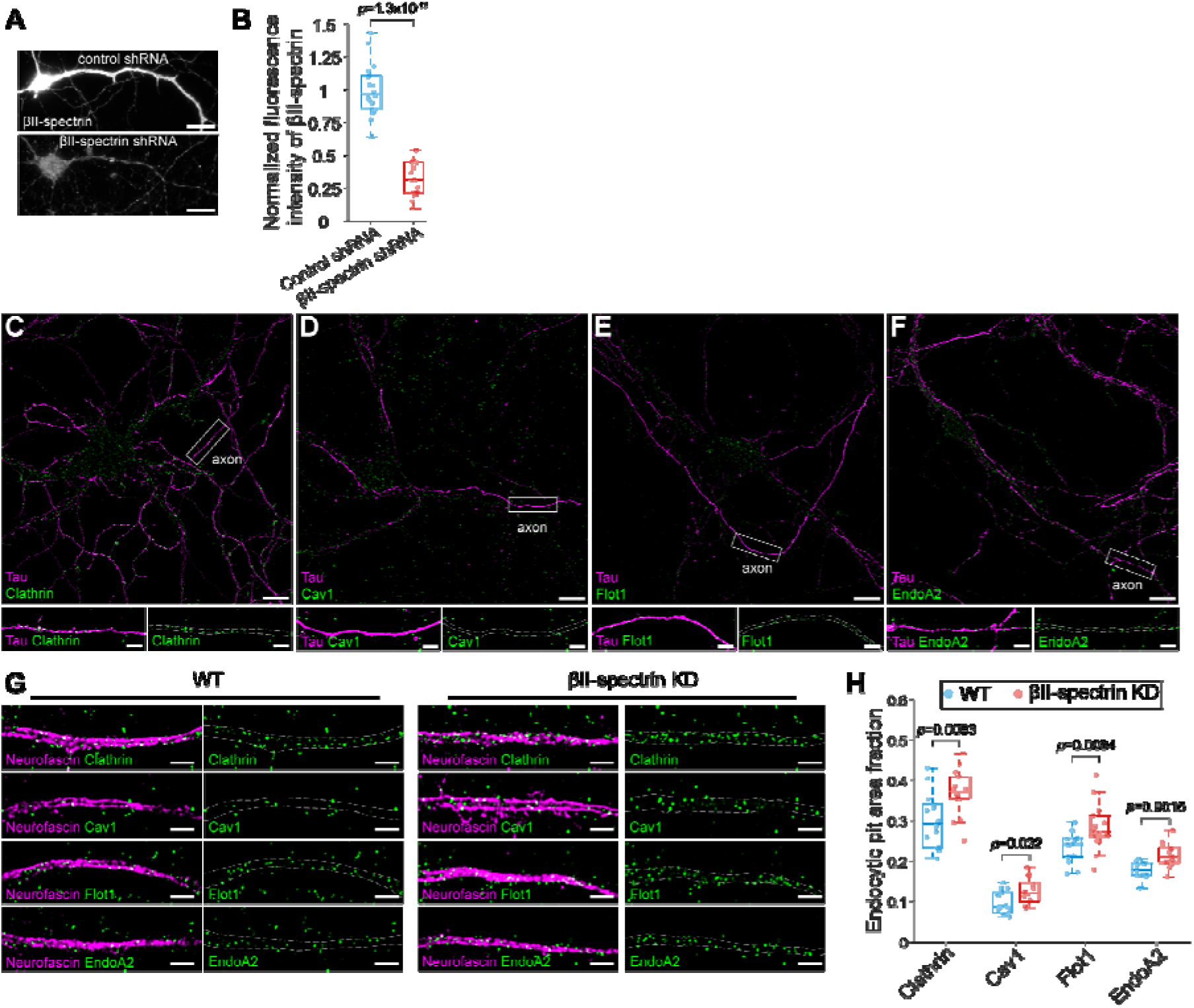
Basal endocytosis activities in distal axons and the effect of MPS disruption on the basal endocytosis in the Axon Initial Segment (AIS), related to Fig. 1. (**A**) Widefield epi-fluorescence images of βII-spectrin in neurons transduced with adenoviruses expressing either scrambled (control) shRNA or βII-spectrin shRNA. Scale bar: 25 µm. (**B**) Box plot showing the normalized fluorescence intensity of βII-spectrin. (**C-F**) Stitched SIM images showing the distributions of endogenous endocytic pits, clathrin (**C**), Cav1(**D**), Flot1 (**E**), or EndoA2 (**F**) in wild-type (WT) neurons. Endocytic pits are shown in green, tau, a distal axon marker, is shown in magenta. Scale bar: 10 µm. Bottom: Enlarged SIM images of the boxed regions on the top, corresponding to distal axon compartments. Scale bar: 2 µm. (**G)** Left: SIM images of neurofascin (magenta) and endogenous endocytic pits (green; clathrin, Cav1, Flot1, or EndoA2) at the AIS of WT neurons. Right: Same as Left, but in βII-spectrin knockdown (KD) neurons. Scale bars: 2 µm. (**H**) Box plot showing the area fraction of endogenous endocytic pits at the AIS of WT and βII-spectrin KD neurons. Boxplots show the median and boundaries (first and third quartile), and the whiskers denote 1.5 times the interquartile range of the box. p-values calculated with two-sided unpaired Student’s t-test.

**Fig. S3.**
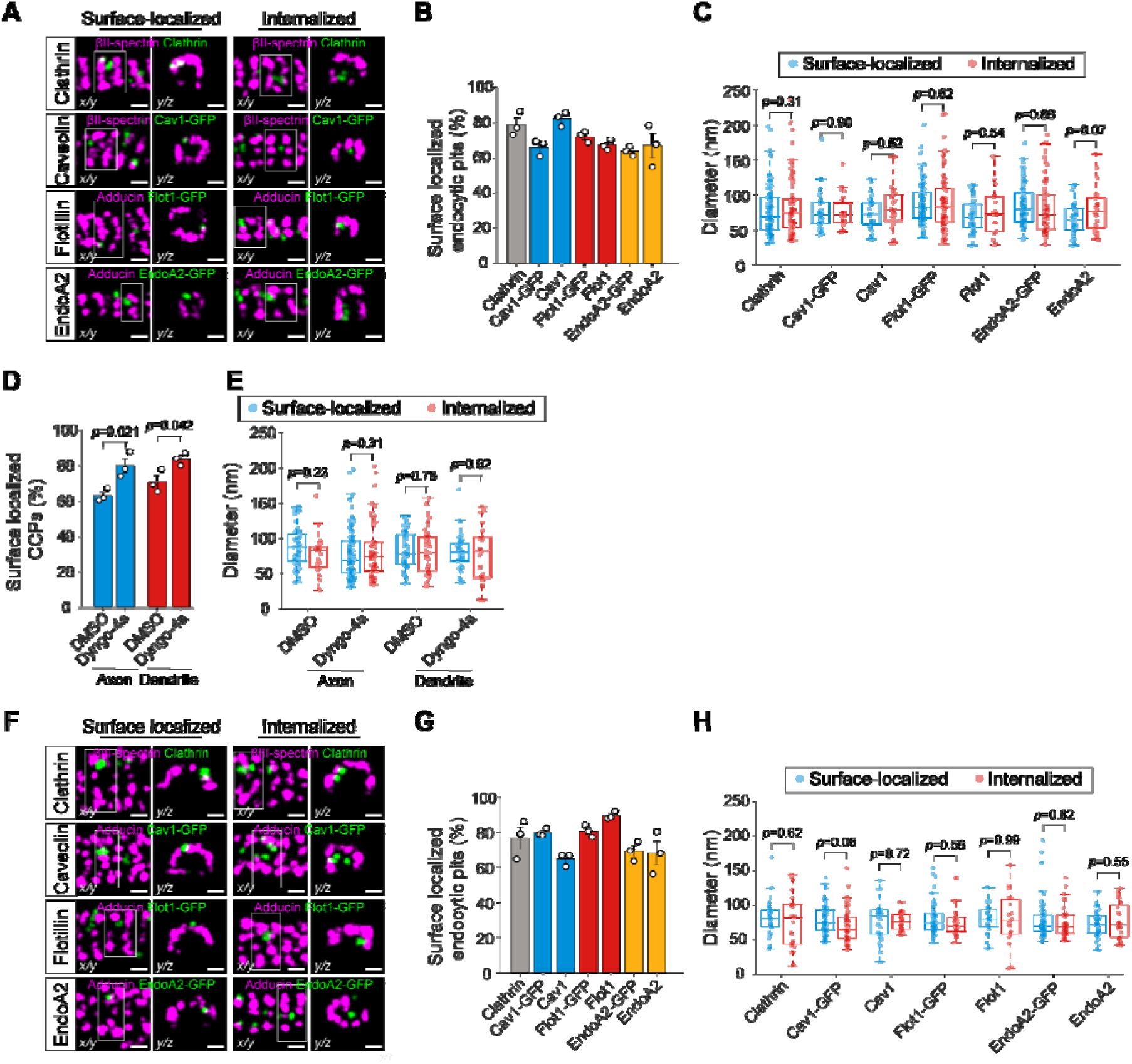
Quantification of surface-localized and internalized endocytic pits in axons and dendrites, related to Fig. 2 and Fig. 3. (**A**) Left: Representative dual-color STORM images of the MPS (magenta) and surface-localized endocytic pits (green) in axons. Endogenous clathrin and exogenously expressed Cav1 were co-stained with βII-spectrin, while exogenously expressed Flot1 and exogenously expressed EndoA2 were co-stained with adducin. The y/z view corresponds to the white-boxed region in the x/y view. Right: Same as Left, but for endocytic pits internalized within the axonal cytoplasm. Scale bars: 200 nm. **(B**) Averaged percentage of surface-localized endocytic pits in axons. Quantification includes only endogenous clathrin, and both endogenous and exogenously expressed Cav1, Flot1, and EndoA2. (**C**) Box plot showing the diameters of surface-localized and internalized endocytic pits in axons. Quantification includes the same categories as in (**B**). (**D**) Averaged percentage of surface-localized CCPs in axons and dendrites under DMSO (control) or dyngo-4a treatment. (**E)** Box plot showing the diameters of membrane-localized and internalized CPPs in axons and dendrites under DMSO or dyngo-4a treatment. (**F-H)** Same as (**A-C)**, but for dendrites. Data are presented as mean ± s.e.m. (n = 3 biological replicates per condition). Boxplots show the median and boundaries (first and third quartile), and the whiskers denote 1.5 times the interquartile range of the box. p-values calculated with two-sided unpaired Student’s t-test.

**Fig. S4.**
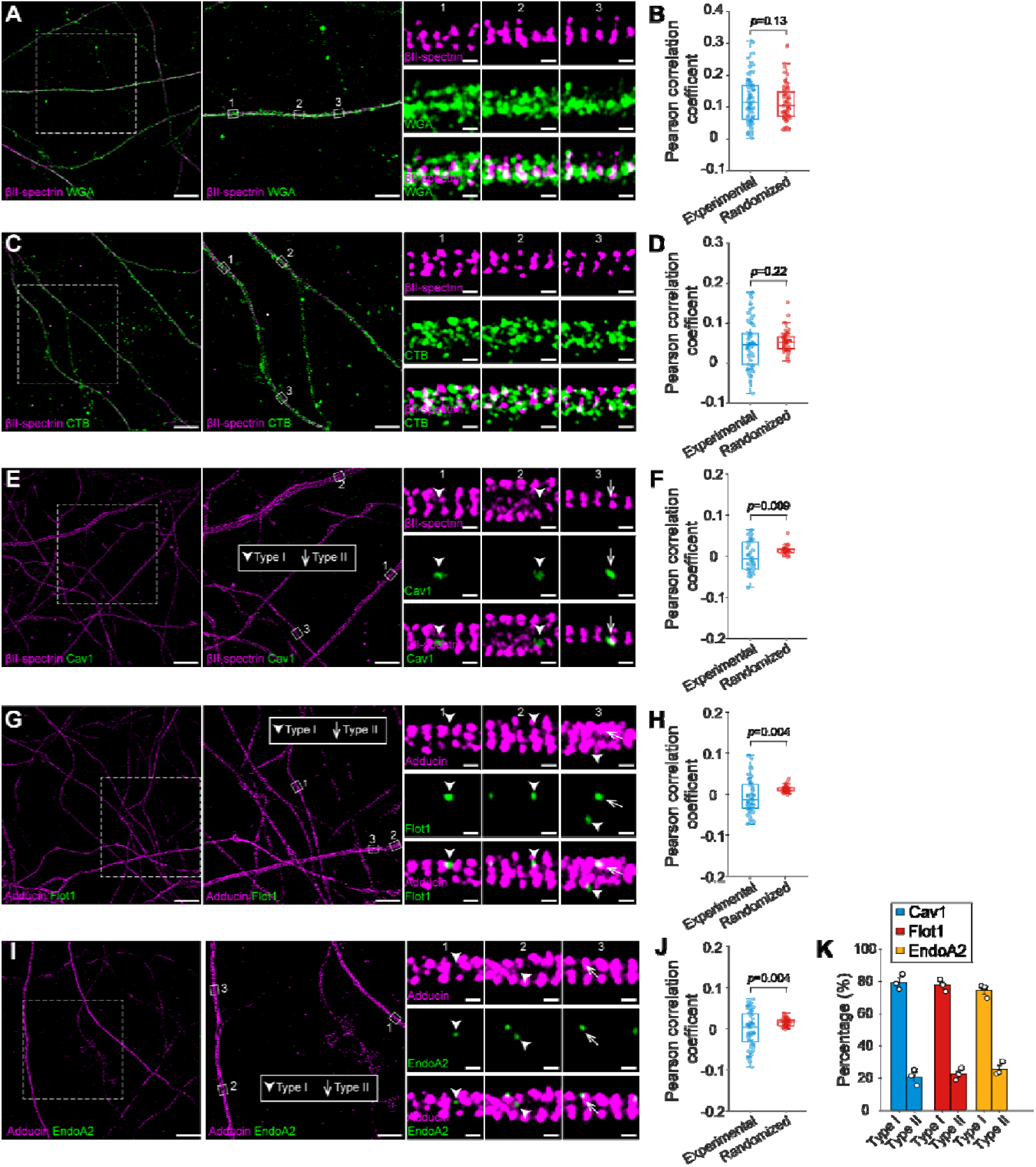
Endogenous endocytic pits are localized within “clearings” of the MPS lattice in axons, related to Fig. 2. (**A**) Dual-color STORM images of βII-spectrin (magenta) and wheat germ agglutinin (WGA, green) in axons. Scale bars: 10 µm (left), 5 µm (middle), 200 nm (right). (**B**) Pearson correlation coefficients between βII-spectrin and WGA under experimental and randomized conditions. (**C, D**) Same as (**A, B**), but for βII-spectrin (magenta) and cholera toxin subunit B (CTB, green). (**E**) Left: Dual-color STORM images of βII-spectrin (magenta) and endogenous Cav1 (green) in axons. Right: Representative enlarged images of Type I and Type II Cav1-pits in the boxed regions. Scale bars: 10 µm (left), 5 µm (middle), 200 nm (right). (**F**) Pearson correlation coefficients between βII-spectrin and endogenous Cav1 under experimental and randomized conditions. (**G, H**) Same as (**E, F**), but for adducin (magenta) and endogenous Flot 1 (green). **(I, J**) Same as (**E, F**), but for adducin (magenta) and endogenous EndoA2 (green). (**K)** Percentages of Type I and Type II pits for endogenous Cav1, endogenous Flot1 and endogenous EndoA2 in axons. Data are presented as mean ± s.e.m. (n = 3 biological replicates per condition). Boxplots show the median and boundaries (first and third quartile), and the whiskers denote 1.5 times the interquartile range of the box. p-values calculated with two-sided paired Student’s t-test.

**Fig. S5.**
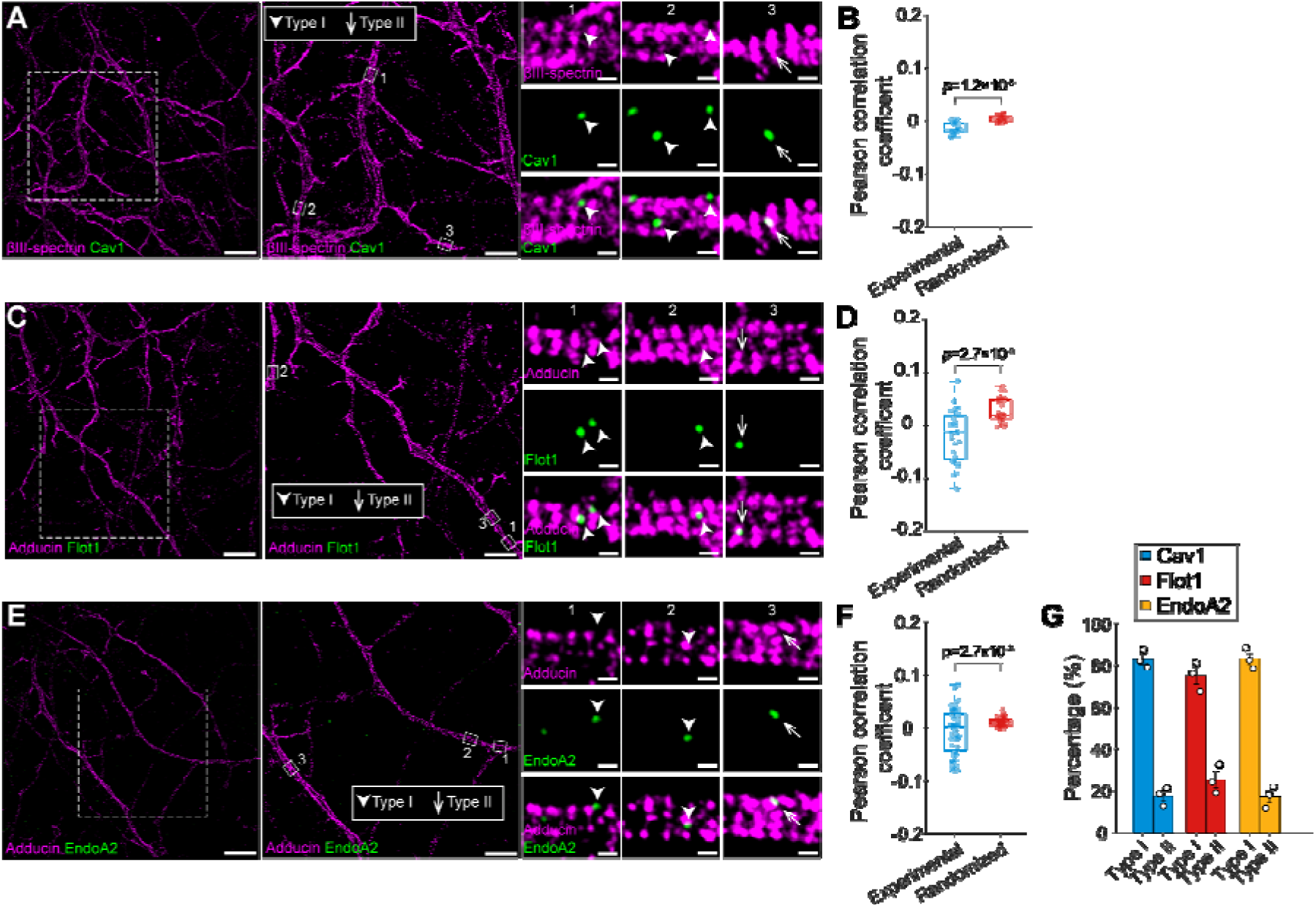
Endogenous endocytic pits are localized within “clearings” of the MPS lattice in dendrites, related to Fig. 3. (**A**) Left: Dual-color STORM images of βIII-spectrin (magenta) and endogenous Cav1 (green) in dendrites. Right: Representative enlarged images of Type I and Type II Cav1-pits from the boxed regions. Scale bars: 10 µm (left), 5 µm (middle), 200 nm (right). (**B**) Pearson correlation coefficients between βIII-spectrin and endogenous Cav1 under experimental and randomized conditions. (**C, D**) Same as in (**A, B**), but for adducin (magenta) and endogenous Flot1 (green). (**E, F**) Same as in (**A, B**), but for adducin (magenta) and endogenous EndoA2 (green). (**G)** Percentages of Type I and Type II pits for endogenous Cav1, endogenous Flot1 and endogenous EndoA2 in dendrites. Data are presented as mean ± s.e.m. (n = 3 biological replicates per condition). Boxplots show the median and boundaries (first and third quartile), and the whiskers denote 1.5 times the interquartile range of the box. p-values calculated with two-sided paired Student’s t-test.

**Fig. S6.**
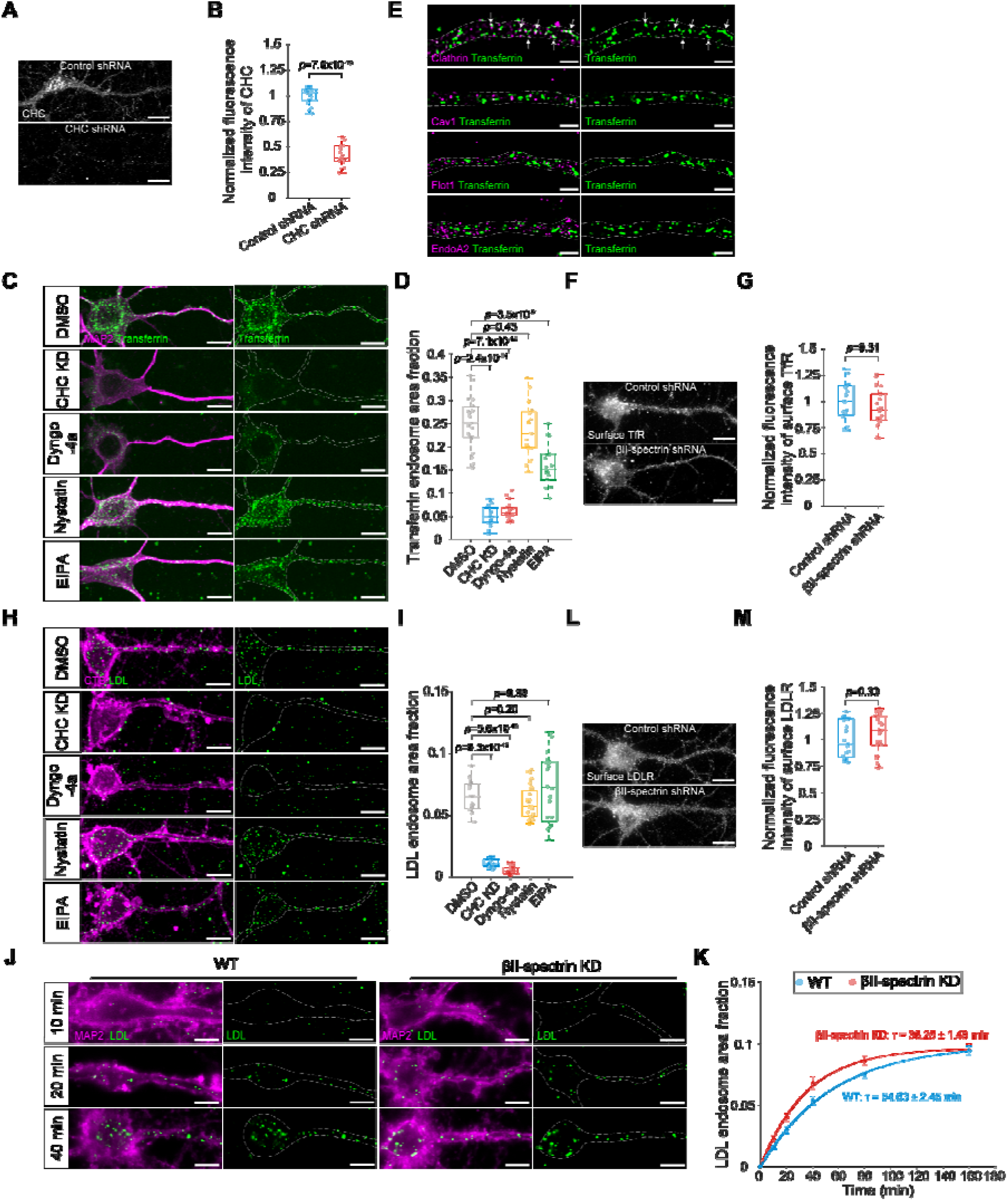
Endocytic pathways for transferrin and LDL endocytosis and the effect of MPS disruption on LDL endocytosis, related to Fig. 4. (**A**) Widefield epi-fluorescence images of clathrin heavy chain (CHC) in neurons transduced with adenoviruses expressing either scrambled (control) shRNA or CHC shRNA. Scale bars: 25 µm. (**B**) Box plot showing the normalized fluorescence intensity of CHC. (**C**) Confocal fluorescence images of MAP2 (magenta) and internalized CF568-transferrin (green) in somatodendritic region of CHC KD neurons pretreated with DMSO followed by CF568-transferrin treatment for 40 minutes, and WT neurons pretreated with DMSO, dyngo-4a, nystatin or EIPA for 30 minutes, followed by CF568-transferrin treatment for 40 minutes. Scale bars: 10 µm. (**D**) Box plot of CF568-transferrin endocytosis in somatodendritic regions of neurons, quantified by the area fraction of transferrin endosomes. (**E**) Dual-color SIM images of internalized CF568-transferin (green) and endocytic pits (magenta; clathrin, Cav1, Flot1 or EndoA2) in dendrites of WT neurons treated with CF568-transferrin for 30 minutes. Regions of strong colocalization are indicated by white arrows. Scale bars: 2 µm. (**F**) Widefield epi-fluorescence images of surface transferrin receptor (TfR) in WT and βII-spectrin KD neurons. Scale bars: 25 µm. (**G)** Box plot showing the normalized fluorescence intensity of surface TfR. (**H**) Widefield epi-fluorescence images of CTB (magenta) and internalized Dil-LDL (green) in somatodendritic region of CHC KD neurons pretreated with DMSO, followed by treatment with Dil-LDL for 40 minutes, and WT neurons pretreated with DMSO, dyngo-4a, nystatin or EIPA for 30 minutes, followed by treatment with Dil-LDL for 40 minutes. Scale bars: 10 µm. (**I**) Box plot of Dil-LDL endocytosis in somatodendritic regions of neurons, quantified by the area fraction of LDL endosomes. (**J**) Left: Widefield epi-fluorescence images of internalized Dil-LDL in somatodendritic region of WT neurons treated with Dil-LDL for 10, 20 and 40 minutes. Right: Same as Left, but in βII-spectrin KD neurons. Scale bars: 10 µm. (**K**) Time course of LDL endocytosis in the somatodendritic regions of WT and βII-spectrin KD neurons, quantified by the area fraction of LDL endosomes. Solid lines represent single-exponential fits. Data are presented as mean ± s.e.m. (**L**) Widefield epi-fluorescence images of surface LDLR in WT and βII-spectrin KD neurons. Scale bars: 25 µm. (**M**) Box plot showing the normalized fluorescence intensity of surface LDLR. Boxplots show the median and boundaries (first and third quartile), and the whiskers denote 1.5 times the interquartile range of the box. p-values calculated with two-sided unpaired Student’s t-test.

**Fig. S7.**
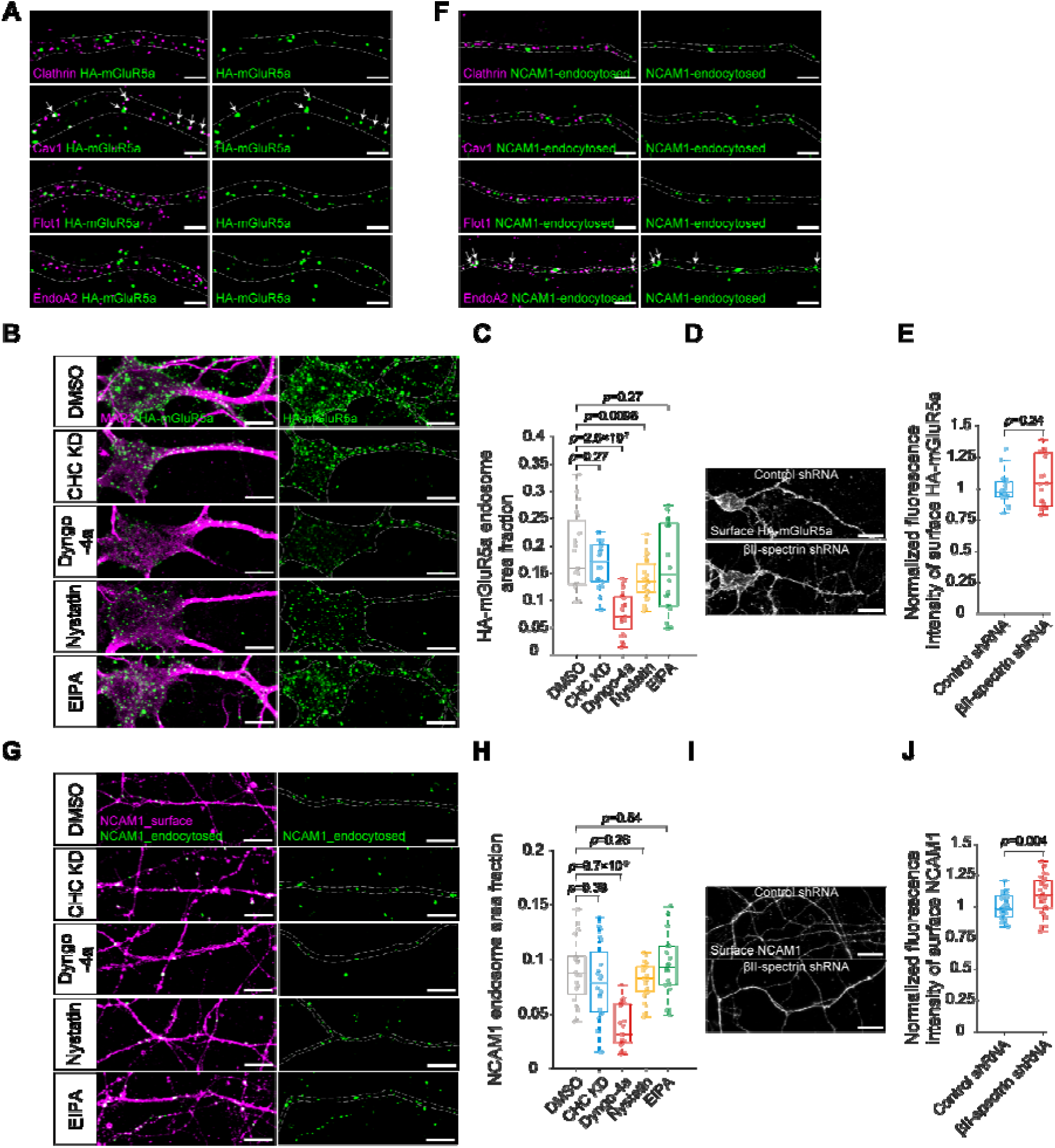
Endocytic pathways for HA-mGluR5a and NCAM1 endocytosis, related to Fig. 4. (**A**) Dual-color SIM images of internalized HA-mGluR5a (green) and endocytic pits (magenta; clathrin, Cav1, Flot1 or EndoA2) in dendrites of WT neurons treated with anti-HA antibody for 30 minutes. Regions of strong colocalization are indicated by white arrows. Scale bars: 2 µm. (**B**) Confocal fluorescence images of MAP2 (magenta) and internalized HA-mGluR5a (green) in the somatodendritic regions of CHC KD neurons overexpressing HA-mGluR5a pretreated with DMSO, followed by treatment with anti-HA antibody for 40 minutes, and WT neurons overexpressing HA-mGluR5a pretreated with DMSO, dyngo-4a, nystatin or EIPA for 30 minutes, followed by treatment with anti-HA antibody for 40 minutes. Scale bars: 10 µm. (**C**) Box plot of HA-mGluR5a endocytosis in somatodendritic regions of neurons, quantified by the area fraction of HA-mGluR5a endosomes. (**D**) Widefield epi-fluorescence images of surface HA-mGluR5a immunostained with anti-HA antibody in WT and βII-spectrin KD neurons. Scale bars: 25 µm.(**E**) Box plot showing the normalized fluorescence intensity of surface HA-mGluR5a. (**F**) Dual-color SIM images of internalized NCAM1 (green) and endocytic pits (magenta; clathrin, Cav1, Flot1 or EndoA2) in axons of WT neurons treated with anti-NCAM1 antibody for 30 minutes. Regions of strong colocalization are indicated by white arrows. Scale bars, 2 µm. (**G**) Confocal fluorescence images of surface NCAM1 and internalized NCAM1 in CHC KD neurons pretreated with DMSO, followed by treatment with anti-NCAM1 antibody for 40 minutes, and WT neurons pretreated with DMSO, dyngo-4a, nystatin or EIPA for 30 minutes, followed by treatment with anti-NCAM1 antibody for 40 minutes. Scale bars: 10 µm. (**H**) Box plot of NCAM1 endocytosis in neurons, quantified by the area fraction of NCAM1 endosomes. (**I**) Widefield epi-fluorescence images of surface NCAM1 immunostained with anti-NCAM1 antibody in WT and βII-spectrin KD neurons. Scale bars: 15 µm. (**J**) Box plot showing the normalized fluorescence intensity of surface NCAM1. Boxplots show the median and boundaries (first and third quartile), and the whiskers denote 1.5 times the interquartile range of the box. p-values calculated with two-sided unpaired Student’s t-test.

**Fig. S8.**
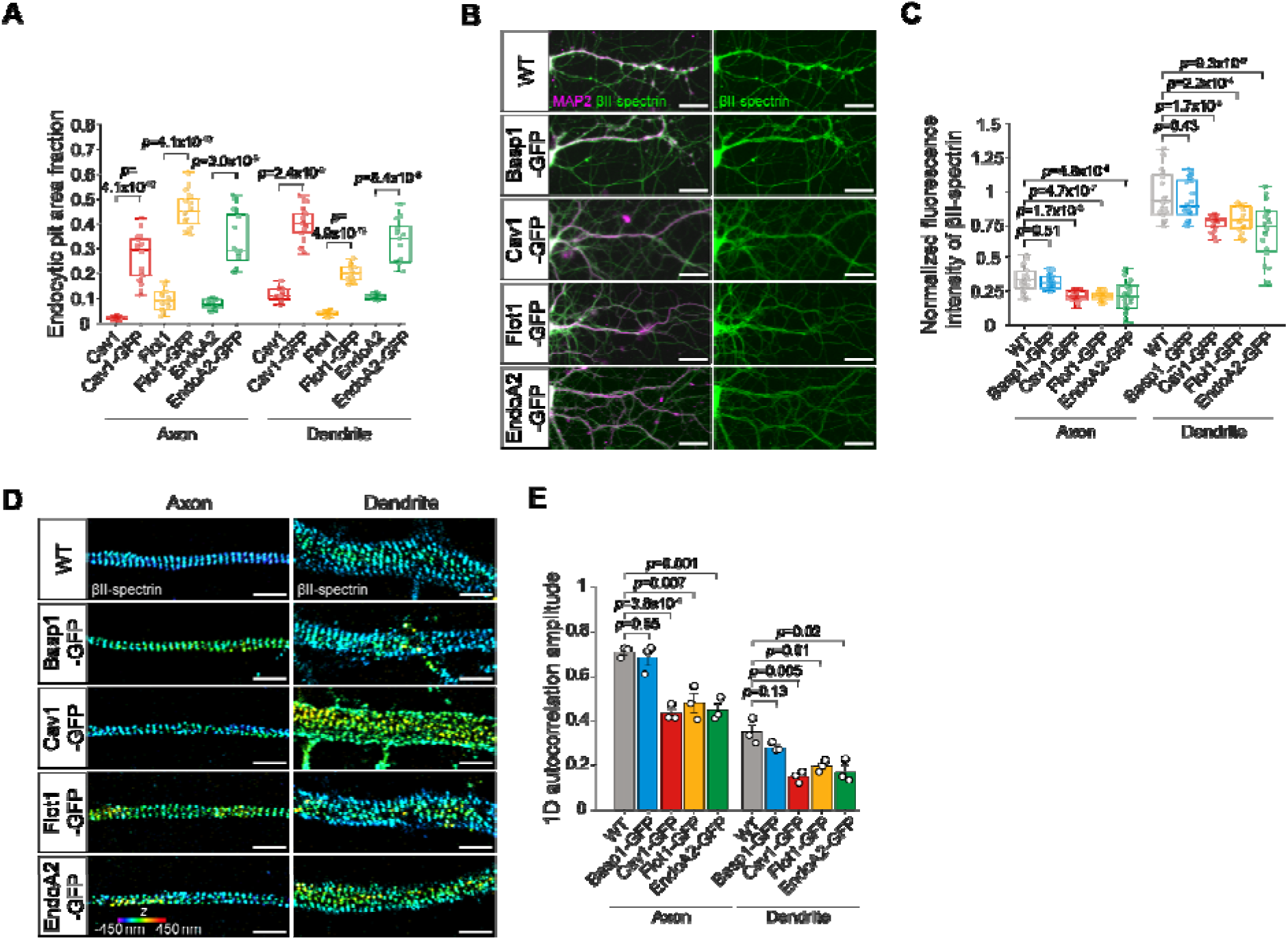
Overexpression of endocytic proteins partially degrades the MPS in neurons, related to Fig. 5. (**A**) Box plot of the area fraction of endogenous and overexpressed endocytic pits in axonal and dendritic neuronal compartments, quantified from the STORM data shown in Figs. 2, and 3, and Extended Data Figs. 4 and 5. (**B**) Widefield epi-fluorescence images of MAP2 (magenta) and βII-spectrin (green) in WT neurons, and neurons overexpressing Basp1, Cav1, Flot1, and EndoA2. Scale bars, 25 µm. (**C**) Box plot of normalized fluorescence intensity of βII-spectrin in axons and dendrites of neurons. (**D**) 3D STORM images of immunostained βII-spectrin in axons and dendrites of WT neurons, and neurons overexpressing Basp1, Cav1, Flot1, and EndoA2. Scale bars, 1 µm. Color scale bar represents the z-coordinate information. (**E**) Averaged 1D autocorrelation amplitude of βII-spectrin in axons and dendrites of neurons. Boxplots show the median and boundaries (first and third quartile), and the whiskers denote 1.5 times the interquartile range of the box. p-values calculated with two-sided unpaired Student’s t-test.

**Fig. S9.**
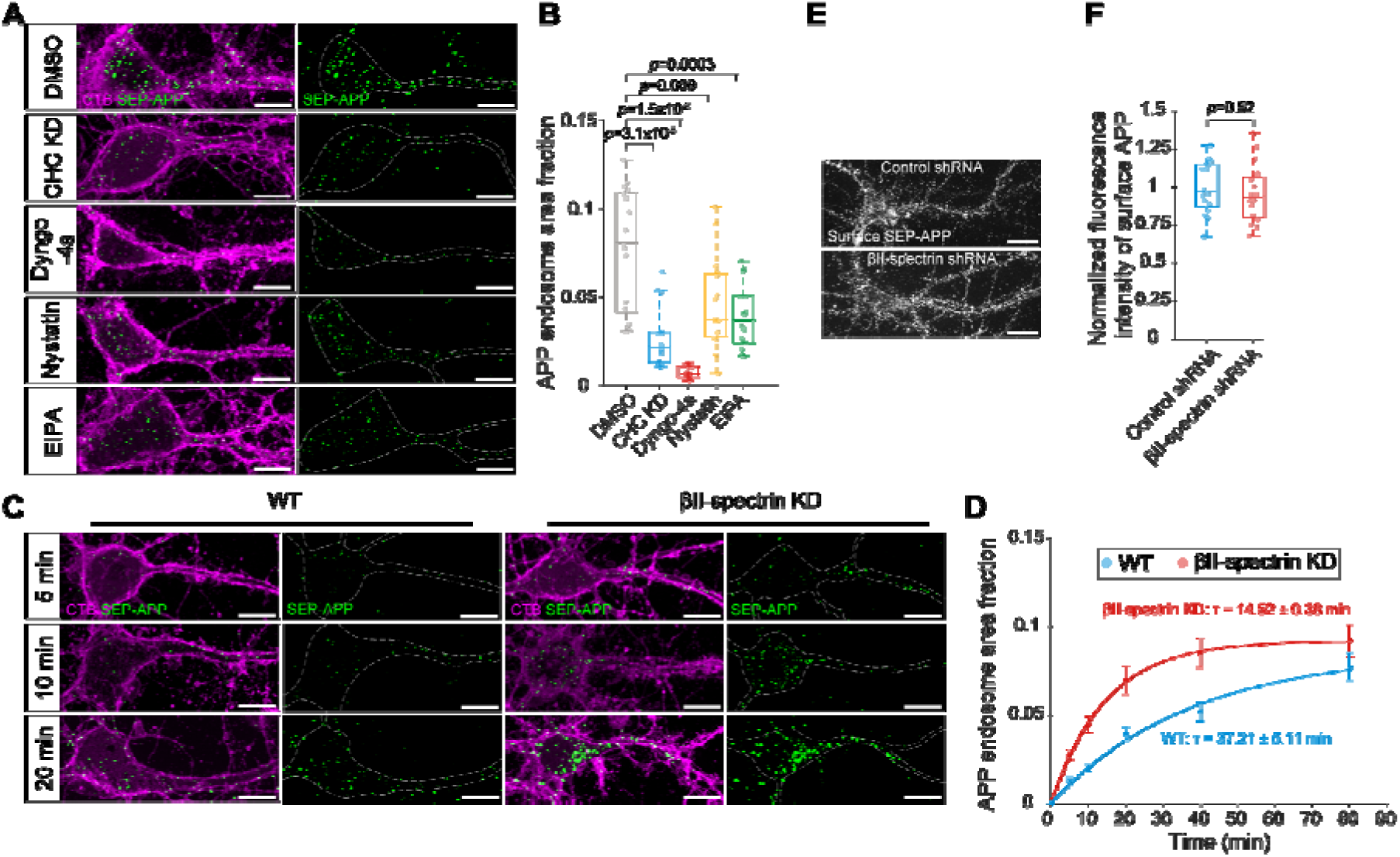
Endocytic pathways of APP endocytosis, related to Fig. 6. (**A**) Confocal fluorescence images of CTB (magenta) and internalized SEP-APP (green) in somatodendritic regions of CHC KD neurons overexpressing SEP-APP pretreated with DMSO, followed by GFP nanobody treatment for 40 minutes, and WT neurons overexpressing SEP-APP pretreated with DMSO, dyngo-4a, nystatin or EIPA for 30 minutes, followed by GFP nanobody treatment for 40 minutes. Scale bars: 10 µm. (**B**) Box plot of SEP-APP endocytosis in somatodendritic regions of neurons, quantified by the area fraction of SEP-APP endosomes. (**C**) Left: Confocal fluorescence images of CTB (magenta) and internalized SEP-APP in somatodendritic region of WT neurons overexpressing SEP-APP treated with GFP nanobody for 5, 10, and 20 minutes. Right: Same as Left, but in βII-spectrin KD neurons. Scale bars: 10 µm. (**D**) Time course of APP endocytosis in the somatodendritic regions of WT and βII-spectrin KD neurons, quantified by the area fraction of APP endosomes. Solid lines represent single-exponential fits to the data. Data are presented as mean ± s.e.m. (**E**) Widefield epi-fluorescence images of surface SEP-APP immunostained with anti-GFP antibody in WT and βII-spectrin KD neurons. Scale bars: 25 µm. (**F**) Box plot showing the normalized fluorescence intensity of surface SEP-APP. Boxplots show the median and boundaries (first and third quartile), and the whiskers denote 1.5 times the interquartile range of the box. p-values calculated with two-sided unpaired Student’s t-test.

